# Associations Between Nutrition, Gut Microbiome, and Health in A Novel Nonhuman Primate Model

**DOI:** 10.1101/177295

**Authors:** Jonathan B. Clayton, Gabriel A. Al-Ghalith, Ha Thang Long, Bui Van Tuan, Francis Cabana, Hu Huang, Pajau Vangay, Tonya Ward, Vo Van Minh, Nguyen Ai Tam, Nguyen Tat Dat, Dominic A. Travis, Michael P. Murtaugh, Herbert Covert, Kenneth E. Glander, Tilo Nadler, Barbara Toddes, John C.M. Sha, Randall S. Singer, Dan Knights, Timothy J. Johnson

## Abstract

Red-shanked doucs (*Pygathrix nemaeus*) are endangered, foregut-fermenting colobine primates which are difficult to maintain in captivity. There are critical gaps in our understanding of their natural dietary habits including consumption of leaves, unripe fruit, flowers, seeds, and other plant parts. There is also a lack of understanding of enteric adaptations, including their unique microflora. To address these knowledge gaps, we used the douc as a model to study relationships between gastrointestinal microbial community structure, diet, and health. We analyzed published fecal samples as well as detailed dietary history from doucs with four distinct lifestyles (wild, semi-wild, semi-captive, and captive) and determined gastrointestinal bacterial microbiome composition using 16S rRNA sequencing. A clear gradient of microbiome composition was revealed along an axis of natural lifestyle disruption, including significant associations with diet, health, biodiversity, and microbial function. We identified potential microbial biomarkers of douc dysbiosis, including *Bacteroides* and *Prevotella*. Our results suggest a gradient-like shift in captivity causes an attendant shift to severe gut dysbiosis, thereby resulting in gastrointestinal issues.

## INTRODUCTION

The primate gastrointestinal (GI) tract is home to trillions of bacteria that play major roles in digestion and metabolism, immune system development, pathogen resistance, and other important aspects of host health and behavior. While there has been substantial progress in understanding the role microbial communities play in human health and disease, less attention has been given to host-associated microbiomes and diet in nonhuman primates (NHPs). Developing a better understanding of the link between primate microbial communities, diet, and health is important not only in the context of primate ecology, but also may have profound implications for use of NHPs as model systems for health and microbial associations.

One colobine primate species, the red-shanked douc (i.e., douc), is of particular interest as a model organism. Because it performs both foregut fermentation and hindgut digestion (Chivers 1994, Lambert 1998), the douc shares digestive characteristics with both humans and ruminant livestock. From a conservation standpoint, it is endangered and fails to thrive in captivity (Agoramoorthy, Alagappasamy, and Hsu 2004, Nijboer 2006, Power, Toddes, and Koutsos 2012). From a health standpoint, this failure to thrive stems foremost from severe gastrointestinal disease, which has been shown in other model organisms and humans to be directly associated with the gut microbiome (Bauchop and Martucci 1968, Edwards, Crissey, and Oftedal 1997, Ensley et al. 1982, Frank et al. 2007, Ley et al. 2006, Overskei et al. 1994, Zhang et al. 2010). Further, in contrast to human studies, the genetic background and dietary profiles of NHPs can be easily and directly ascertained or manipulated, which is critical as both dietary and genetic factors have been implicated in modulation of the microbiome. The douc thus represents a model organism whose relevance may span domains of conservation and microbial ecology, and inform both human and livestock health.

Here, by comparing four different populations of the same species along a captivity/wildness (i.e., lifestyle) gradient, we sought to determine whether such a gradient in lifestyle also manifests in a gradient of gut microbiota and diet, and to assess whether any such trends corroborate health status. Furthermore, examining one species living four different lifestyles allows one to examine the influence of environment independent of interspecific host variation on shaping gut microbial community structure. As microbes may act as indicators for health of the host (Frank et al. 2007), these results may allow for the development of predictive biomarkers to improve NHP health and management. As some microbial trends hold across species boundaries in other model systems (Anderson et al. 2012, Kohl, Skopec, and Dearing 2014, Roeselers et al. 2011, Wang et al. 2013, Zhao et al. 2013), some biomarkers may translate to human and ruminant health as well.

Here, we focus on a subset of a rich dataset we collected over a two-year period across three countries, including a comprehensive sampling of the majority of doucs in captivity. Some of these data have been published previously (Clayton et al. 2016) in a broader meta-analysis framework, where a putative convergence was observed across various primate species (including humans) with increasing levels of generalized lifestyle disruption. While the broadness of such an overview was valuable in demonstrating overall trends across species, it may also mask intraspecific effects underlying a lifestyle gradient, and also limits the resolution and interpretability of correlations that can be drawn to specific dietary, health, and lifestyle components, which themselves may play very different biological roles in the various species under investigation. By focusing in depth on the dietary and microbial facets associated with a single species across increasingly unnatural lifestyle conditions, we are powered to make specific conclusions relating these covariates in a common context.

## METHODS

### Study site, subjects, and sample material

Fecal samples (n = 111) were collected opportunistically immediately after defecation from captive (n = 12), semi-captive (n = 15), semi-wild (n = 18), and wild (n = 66) red-shanked doucs (*Pygathrix nemaeus*) in 2012-2013. The microbiome samples were analyzed previously in a meta-analysis examining broader relationships between captivity and the microbiome (Clayton et al. 2016). Two captive, seven semi-captive, and eighteen semi-wild red-shanked doucs were sampled. Fecal samples (n = 26) were collected from seven known wild individuals. Remaining fecal samples (n = 40) collected from the wild population originated from unknown individuals. Doucs housed at the Endangered Primate Rescue Center (EPRC) in Ninh Binh, Vietnam served as the semi-wild population, as the doucs there live a lifestyle (environment and diet) representing an intermediate state between wild and semi-captive. Specifically, these doucs are fed exclusively plants, and thus are not offered any supplemental dietary items, such as ripe fruits, vegetables, or vitamin supplements, all of which are fed to doucs housed at traditional zoological institutions. Doucs housed at the Singapore Zoo served as the semi-captive population, as they live a lifestyle (environment and diet) representing an intermediate state between semi-wild and captive. Doucs housed at the Philadelphia Zoo served as the captive population, as they live in artificial environments compared to their semi-wild and wild counterparts. Doucs inhabiting Son Tra Nature Reserve, Da Nang, Vietnam (16°06’—16°09’N, 108°13’—108°21’E) served as the wild population in this comparative study (Lippold and Thanh 2008, Ulibarri 2013) (Supplemental Figure 1). Son Tra is located only 10 km from the heart of Da Nang City, which is the third largest city in Vietnam. The nature reserve is comprised of 4,439 total ha and, of those, 4,190 ha is covered by both primary and secondary forests (Lippold and Thanh 2008).

### Genomic DNA extraction

Total DNA from each fecal sample was extracted as described with some modifications (Yu and Morrison 2004). Briefly, two rounds of bead-beating were carried out in the presence of NaCl and sodium dodecyl sulfate, followed by sequential ammonium acetate and isopropanol precipitations; precipitated nucleic acids were treated with DNase-free RNase (Roche); and DNA was purified with the QIAmp^^®^^ DNA Stool Mini Kit (QIAGEN, Valencia, CA), according to manufacturer’s recommendations. DNA quantity was assessed using a NanoDrop 1000 spectrophotometer (Thermo Fisher Scientific Inc, Massachusetts, USA).

### Bacterial 16S rRNA PCR amplification and Illumina MiSeq sequencing

The bacterial 16S rRNA gene was analyzed using primers 515F and 806R, which flanked the V4 hypervariable region of bacterial 16S rRNAs (Caporaso et al. 2012). The oligonucleotide primers included Illumina sequencing adapters at the 5’ ends and template specific sequences at the 3’ ends. The primer sequences were: 515F (forward) 5’ GTGCCAGCMGCCGCGGTAA 3’ and 806R (reverse) 5’ GGACTACHVGGGTWTCTAAT 3’ (Caporaso et al. 2012). The 16S rRNA PCR amplification protocol from the earth microbiome project was used (Gilbert et al. 2010). Each sample was amplified in two replicate 25-μL PCR reactions and pooled into a single volume of 50 μL for each sample. The amplification mix contained 13 μL of PCR grade water (MoBio, Carlsbad, CA), 10 μL of 5 PRIME HotMasterMix (5 PRIME, Gaithersburg, MD), 0.5 μL of each fusion primer, and 1.0 μL of template DNA in a reaction volume of 25 μL. PCR conditions were an initial denaturation at 94°C for 3 m; 35 cycles of 94°C 45 s, 50°C for 60 s, and 72°C for 90 s; and a final 10 m extension at 72°C. Following PCR, concentration of PCR products was determined by a PicoGreen assay. Equal amounts of samples were pooled, and size selection was performed using the Caliper XT (cut at 386 bp +/- 15%). Final quantification was performed via a PicoGreen assay and assessment on a Bioanalyzer 2100 (Agilent, Palo Alto, California) using an Agilent High Sensitivity chip. The PCR amplicons were sequenced at the University of Minnesota Genomics Center (UMGC) using Illumina MiSeq and 2x300 base paired-end reads (Illumina, San Diego, California).

### 16S Data analysis

Raw sequences were analyzed with QIIME 1.8.0 pipeline (Caporaso et al. 2010). The demultiplexed sequences from the UMGC were subjected to the following quality filter: 150 bp < length < 1,000 bp; average quality score > 25. Preprocessed sequences were then clustered at 97% nucleotide sequence similarity level. For the diversity and taxonomic analyses, the open-reference-based OTU picking protocol in QIIME was used with GreenGenes 13_8 as the reference database (DeSantis et al. 2006) using the USEARCH algorithm (Edgar 2010). Unmatched reads against the reference database which also did not cluster later in the open reference pipeline were excluded from the downstream analysis. Read depth was relatively uniform across lifestyles (Supplemental Figure 2). Taxonomy information was then assigned to each sequence cluster using RDP classifier 2.2 (Wang et al. 2007). Closed-reference OTUs of chloroplast origin were filtered out with QIIME, and samples were rarefied to 52,918 reads for the downstream analysis.

For the closed-reference-only analyses, including PICRUSt and chloroplast analyses, the raw FASTQ files were processed with SHI7 (Al-Ghalith et al. 2017), a wrapper script that detected and removed TruSeq v3 adaptors with trimmomatic (Bolger, Lohse, and Usadel 2014), stitched the R1 and R2 reads together with FLASh (Magoč and Salzberg 2011), performed quality trimming from both ends of the stitched reads until a minimum quality score ≥ 32 was reached, and filtered out reads with average quality score < 36. 88.5% of all original sequences were retained after QC, resulting in an average read length of 254 bases and average quality score of 37.6. Closed-reference picking was performed at 95% similarity level with the taxonomy-aware exhaustive optimal alignment program BURST (Al-Ghalith and Knights 2017) against a database of all RefSeq chloroplast sequences in phyla Chlorophyta (green algae) and Streptophyta (land plants) as of 06/27/2017, a total of 1,506 chloroplast reference sequences. The same closed-reference procedure was also used to re-pick OTUs against GreenGenes 13_8 for use with PICRUSt, as the latter is reliant on closed-reference GreenGenes IDs for functional prediction.

Alpha diversity (including chao1, shannon, and simpson diversity metrics) and beta diversity analysis (including Bray-Curtis, weighted and unweighted UniFrac metrics) (Lozupone and Knight 2005), were performed and plotted through a combination of wrapper scripts in QIIME and custom R scripts using the vegan, ape, ggplot2, and phyloseq packages (McMurdie and Holmes 2013, Oksanen et al. 2007, Paradis, Claude, and Strimmer 2004, Wickham 2016). ANOVA was used to assess the statistical significance of alpha diversity variation among population, and Adonis was used to assess whether populations significantly differed by beta diversity (Oksanen et al. 2007).

The functional profiles of the microbial sample were investigated using PICRUSt (Phylogenetic Investigation of Communities by Reconstruction of Unobserved States) (Langille et al. 2013), which predicts Kyoto Encyclopedia of Genes and Genomes (KEGG) module abundances within a microbial community based on 16S rRNA surveys. Within this pipeline, relative abundances of OTUs were normalized by 16S rRNA copy number, after which centered log ratio transformation was applied with detection-limit zero replacement. Metagenomic contents were predicted from the KEGG catalogue (Kanehisa and Goto 2000). The mean Nearest Sequenced Taxon Index (NSTI) for all lifestyles was below 0.18 (Supplemental Figure 3). To assess degree of correlation between each functional module and population group ordered by degree of wildness, polyserial correlation was used with ordered lifestyle groups (captive < semi-captive < semi-wild < wild) via the polycor package (Fox 2007). Statistical significance of this correlation was ascertained by computing a p-value from the polyserial chi-square rho and degrees of freedom, followed by Holm family-wise error rate correction (alpha < 0.05). Additionally, the captive and wild populations were compared pairwise using the non-parametric Wilcoxon Rank-Sum test followed by Holm adjustment. All associations for which absolute rho < 0.3, adj. p > 0.05, or rho confidence p > 0.05 were considered insignificant.

Differential taxon abundance testing was also performed. OTUs were binned additively according to taxonomy at the genus level (or at lowest characterized level if genus was uncharacterized for a given OTU). The resulting taxa were then filtered such that only taxa present in at least 3 samples and with 0.01% average abundance throughout the dataset were retained, leaving 75 distinct taxa at or below genus level. The OTU table was normalized by centered log-ratio with least-squares zero interpolation (Templ, Kowarik, and Filzmoser 2011) to allow for valid compositional covariate testing (Tsilimigras and Fodor 2016). Similarly to the PICRUSt KEGG module differential abundance testing described above, statistical significance of association between each taxon and the sample populations was assessed with polyserial correlation across groups in order of wildness, as well as Wilcoxon rank-sum tests of the two extrema (captive vs wild lifestyles), followed by Holm adjustment. All associations for which absolute rho < 0.3, adj. p > 0.05, or rho confidence p > 0.05 were considered insignificant. For heatmaps, only features with absolute rho > 0.6, adj. p < 0.05, and rho confidence p < 0.05 are displayed for clarity.

### Data deposition

All sequencing data are deposited at the European Bioinformatics Institute under project number PRJEB11414. Additionally, all R code and raw non-sequence data used for these analyses is freely available on the project GitHub site located at https://github.com/jbclayton83/douc-microbes-paper.

### Analysis of diet components

One population of wild doucs was observed between January and August 2013 in Son Tra Nature Reserve, Danang, Vietnam. All occurrences of observed feeding behaviors were recorded. Identified plant parts ingested were recorded and reachable feeding trees were marked. The plant parts of specific trees which were prevalent in their diet and were available in sufficient quantities were sampled and dried to 95% dry matter as per a previously established method (Barnett 1995). Samples were sent to the Biochemical Lab at The Agriculture and Forestry University in Ho Chi Minh City, Vietnam, for chemical analysis. Concentrations of crude protein, simple sugars, crude fiber, calcium, sodium, manganese, potassium and iron were determined on a dry matter basis, all of which follow AOAC methods 920.152, 973.18C, and 974.06 (AOAC 2012). Additionally, all plants fed to semi-wild doucs during a two week period in October 2012 were also sent for chemical analysis for comparison. Chemical compositions of semi-captive and captive diets were also analyzed. Nutrients were compiled from the laboratory results on a concentration per dry matter basis. Wild and semi-wild nutrient contents were constructed by observed frequency. Semi-captive and captive nutrient contents were assembled purely from diets as given to their specimens.

## RESULTS

### Microbiome diversity declines according to lifestyle and habitat disruption

Fecal microbiome diversity showed a steady decline from wild towards captive environments, as we described previously (Clayton et al. 2016). The number of OTUs in the doucs decreased in a gradient-like fashion with the highest number in wild doucs (4231.68 ± 584.37 OTUs), and the lowest number in captive doucs (2332.08 ± 180.30 OTUs). Consistent with the gradient hypothesis, the semi-wild doucs (2845.50 ± 494.98 OTUs) and semi-captive doucs (2696.93 ± 417.00 OTUs) were intermediate. Pairwise comparison of all populations by OTU abundance showed statistically significant differences in OTU count biodiversity between all groups (p < 0.01) (Figure 1a). In addition to investigating overall OTU diversity (i.e., number of OTUs) amongst the four unique douc populations, other alpha diversity (i.e., within-sample diversity) metrics were calculated. The mean Shannon diversity index, which measures species evenness, was highly significant across the four douc populations (wild: 7.86 ± 0.34; semi-wild: 7.07 ± 0.55; semi-captive: 7.11 ± 0.53; captive: 6.65 ± 0.53; ANOVA, p=4.3×10^-18^). Based on the calculated Shannon diversity indices, the wild douc microbiome was the most even of the four douc populations (Figure 1b).

**Figure 1.**
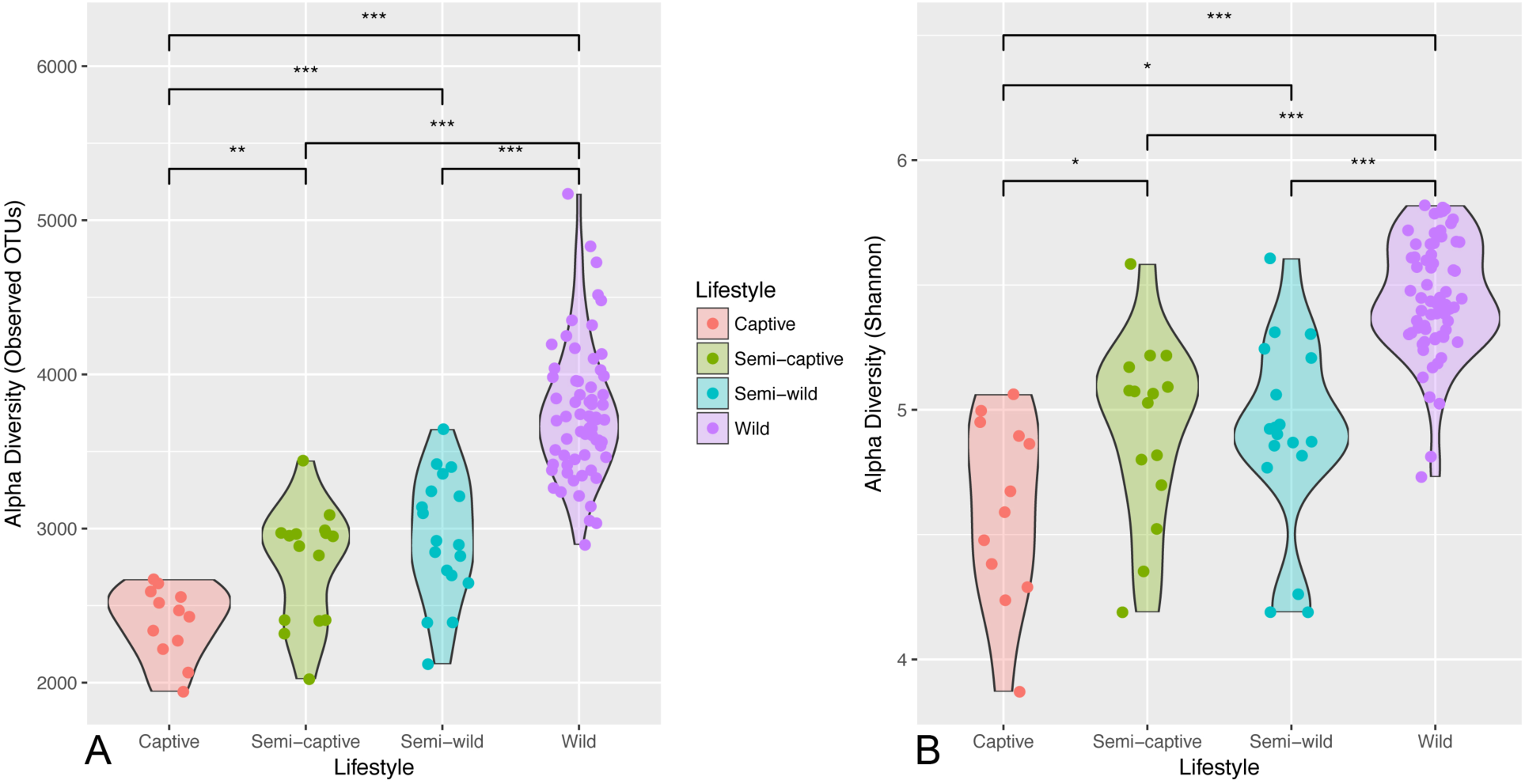
Diminished alpha diversity in red-shanked douc microbiomes across lifestyles. Violin plots of gut microbial alpha diversity across the 4 lifestyles according to (a) the number of species-like operational taxonomic units (OTUs) generated by open-reference OTU picking in the gut microbiome, and (b) the Shannon diversity index. The width of the shape corresponds to the distribution of samples (strips overlaid as strip chart), and asterisks denote significant differences at *p* < 0.05 (*), *p* < 0.01 (**), and *p* < 0.001 (***). Under both metrics, the wild population exhibits the highest biodiversity, which appears to diminish as a gradient with level of captivity to the captive population, which has the lowest.

Beta diversity calculations were performed to assess whether significant differences between populations were present, using unweighted UniFrac distance (Figure 2), as well as the weighted UniFrac and non-phylogenetic Bray-Curtis metrics (Supplemental Figure 4). Analysis of unweighted UniFrac distance measurements is most effective at detecting differences in community membership when considering abundance differences among rare taxa (Chen et al. 2012). An Adonis test on unweighted UniFrac distances revealed that fecal microbiome grouped statistically by douc population (Adonis p = 0.001), suggesting that each douc population had a unique microbiome. It also suggests that lifestyle has a major influence on gut microbial community structure, as doucs living under the most unnatural conditions had gut microbiomes most disparate from free-living wild doucs. Overall, the results of our beta diversity analyses indicated that microbiome composition was distinct for each of the four douc populations examined in this study at the 97% OTU and genus levels. Further, Figure 2 reveals a clear gradient by naturalness of lifestyle along PC1, the primary axis of differentiation.

**Figure 2.**
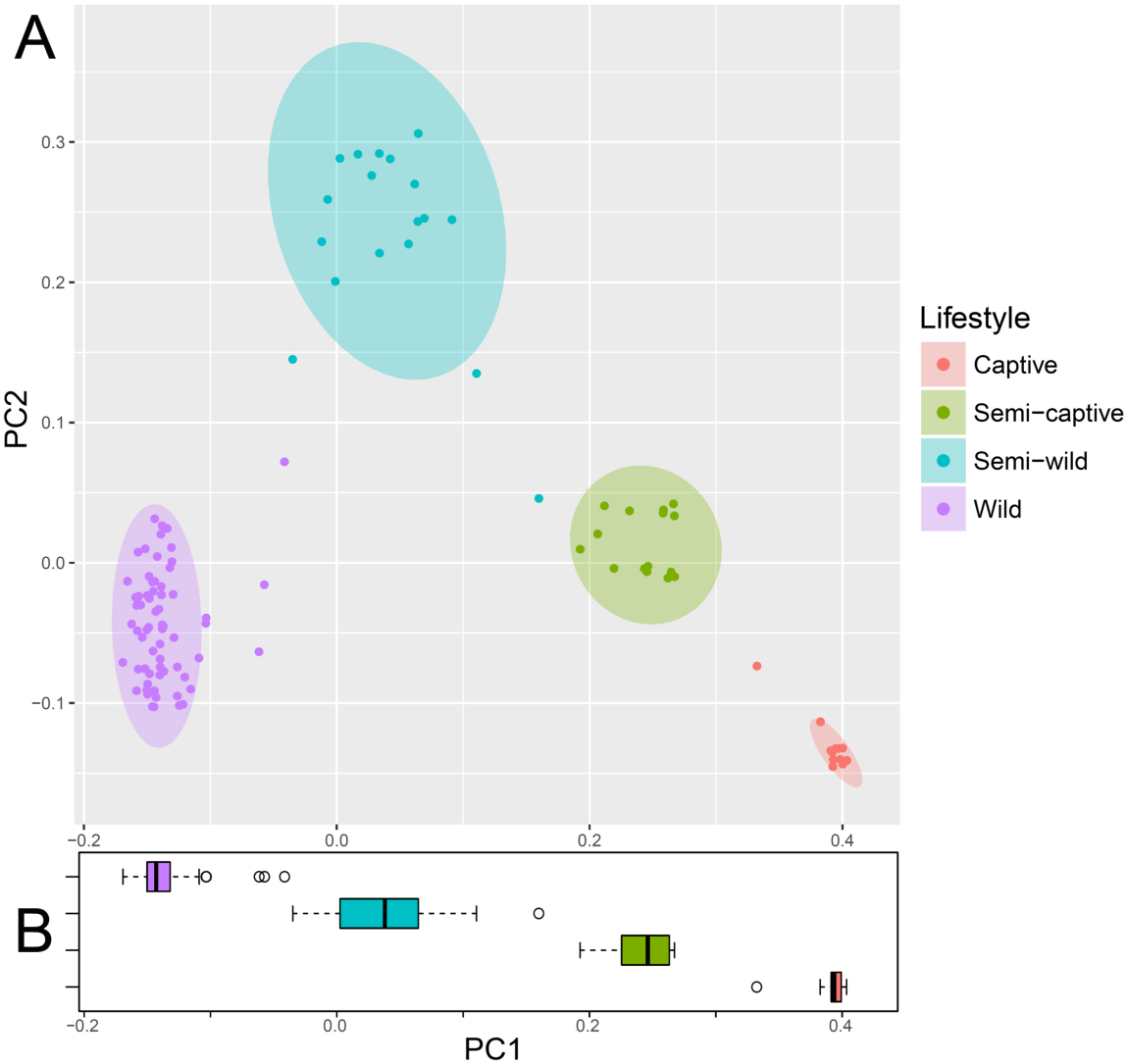
Principal coordinates plot showing (a) unweighted UniFrac ordination and (b) box plot of PC1 by population showing ecological distance between gut microbial communities in wild, semi-wild, semi-captive, and captive red-shanked doucs. All samples were obtained with the same protocol for V4 16S rRNA sequencing, and open-reference OTU picking was used. Douc microbiomes clearly clustered by population suggesting that each douc population had a unique microbiome, and thus were highly distinctive. Lifestyle has a major influence on gut microbial community structure, as doucs living under the most unnatural conditions (captive) had gut microbiomes most disparate from wild doucs (i.e., natural).

### Differential taxonomic abundance analysis by lifestyle

Broad phylum-level taxonomic summarization revealed trends among the fecal microbiomes of the four douc populations included in this study. The fecal microbiomes of wild, semi-wild, semi-captive, and captive doucs were dominated by the phylum Firmicutes. Bacteroidetes was found in very low abundance in both the wild and semi-wild populations. In contrast, Bacteroidetes was the second most abundant phylum found in both the semi-captive and captive populations. Additionally, Verrucomicrobia was much more abundant in the semi-wild fecal microbiome than the other lifestyles examined (Supplemental Figure 5; Supplemental Figure 6). *Bacteroides* and *Prevotella*, as well as *Methanosphaera*, *CF231*, *Treponema*, and *YRC22* were highly positively correlated with captivity level (all polyserial rho ≥ 0.71, p < 1x10^-6^) (Figure 3; Figure 4; Supplemental Figure 6; Supplemental Table 1). Conversely, *Adlercreutzia*, *Anaerostipes*, *Blautia*, *Campylobacter*, *Dehalobacterium*, *Dorea*, and *Oscillospira* were much less abundant with increasing captivity (all polyserial rho ≤ −0.65, p < 1x10^-6^) (Figure 3; Figure 4; Supplemental Figure 6; Supplemental Table 1). Although the genus *Akkermansia* shows a similar trend (polyserial rho = −0.57, p = 1.29x10^-09^), its abundance peaks slightly in the semi-wild population (Figure 3; Figure 4; Supplemental Figure 6).

**Figure 3.**
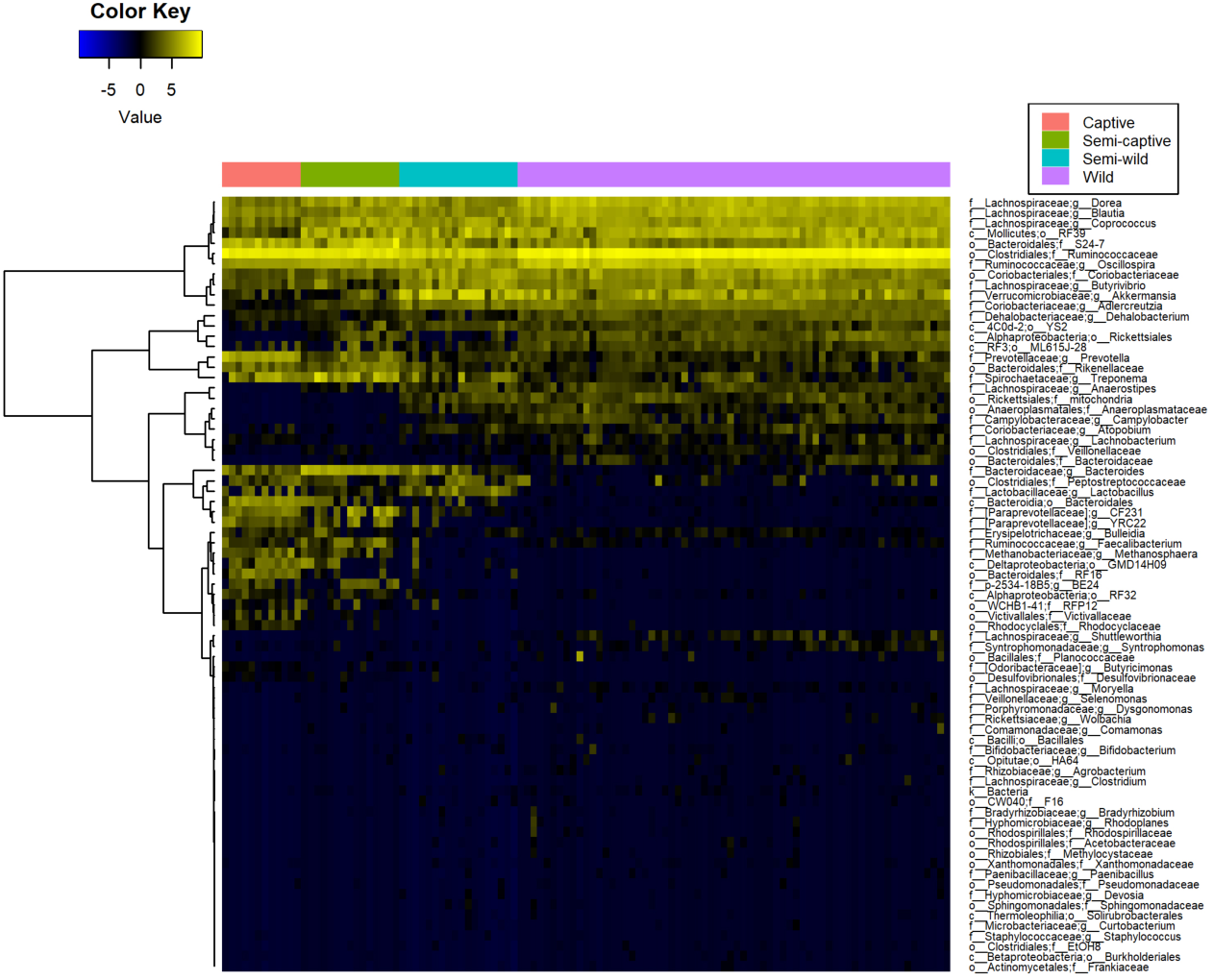
Heatmaps of differentially abundant microbial taxa at the genus level in red-shanked doucs living four distinct lifestyles. Taxa are displayed with polyserial correlations (rho) above 0.3, rho estimate adjusted p < 0.05, and (pairwise) Wilcoxon rank-sum adjusted p-value for wild and captive lifestyles < 0.05. Color represents intensity of centered log ratio abundances along gradient of color scale shown.

**Figure 4.**
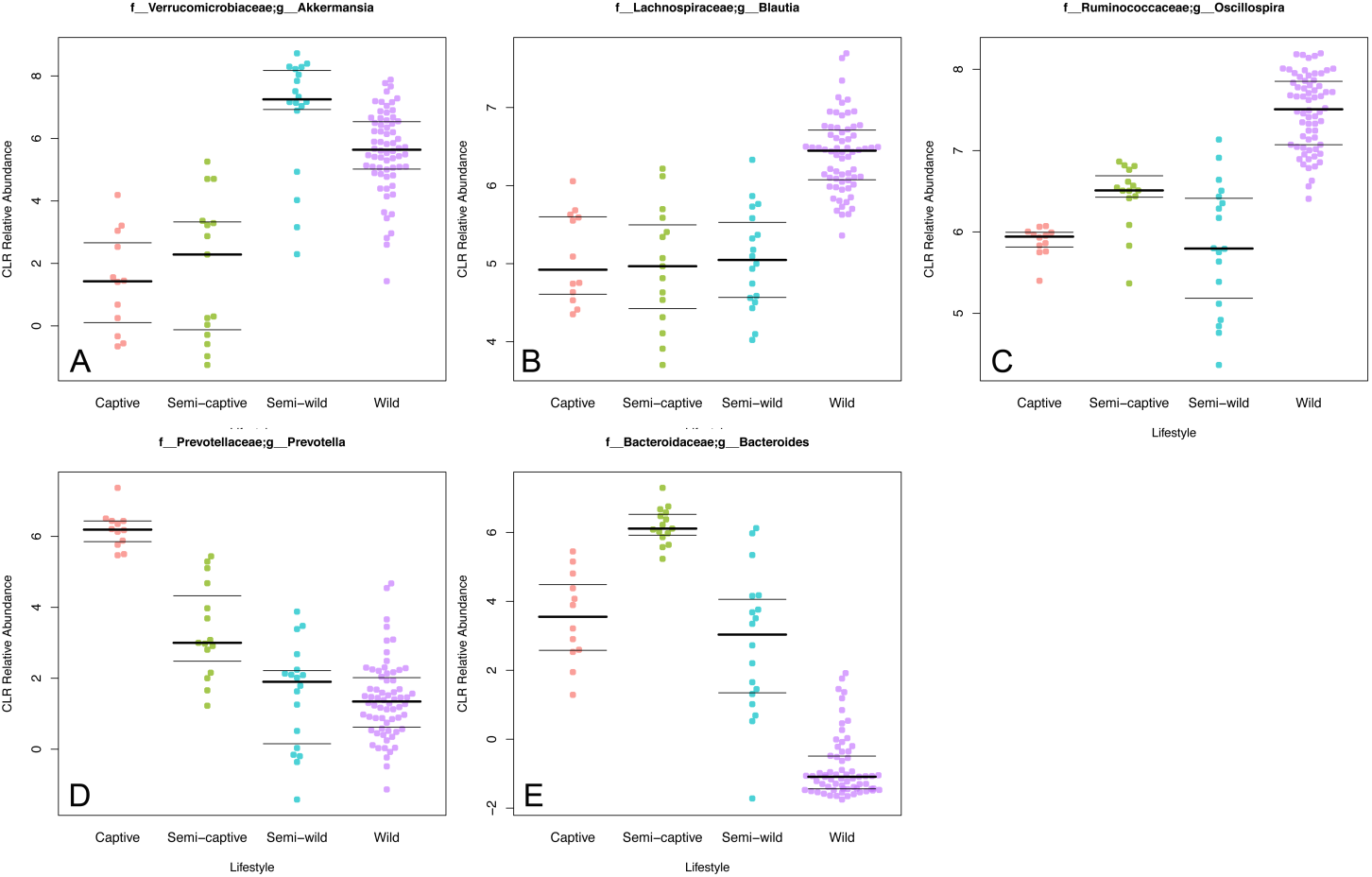
Beeswarm plots displaying gradient-like patterns of selected microbial taxa. A beeswarm plot of the arcsine square root relative abundance of bacterial genera *Bacteroides*, *Prevotella*, *Oscillospira*, *Blautia*, and *Akkermansia* shown in wild, semi-wild, semi-captive, and captive douc populations. All samples were obtained with the same protocol for V4 16S rRNA sequencing, and open-reference OTU picking was used. Red-shanked doucs acquire *Bacteroides* and *Prevotella*, and lose *Oscillospira*, *Blautia*, and *Akkermansia* in captivity. The presence of *Akkermansia* was most associated with a semi-wild lifestyle.

A full-rank, species-level, plant-inclusive heatmap was also generated under more stringent taxa selection criteria (absolute polyserial rho > 0.45, pairwise p < 0.01) but with sample (column) clustering also performed to ascertain whether hierarchical clustering of these taxa abundance profiles alone would be able to recover a similar separation between lifestyles as observed in the Unweighted UniFrac ordination. The unsupervised clustering of these features correctly recovered the group membership of all samples, as observed by the preservation of the sample group labels without gaps or shuffling between lifestyles (Supplemental Figure 7a). With even more stringent selection criteria (polyserial rho > 0.75, pairwise p < 0.001), there is some overlap of samples between neighboring lifestyle groups along the gradient; specifically, it appears as if some members of the “Semi-wild” group have mingled with the adjacent “Semi-captive” and “Wild” groups along the gradient, but the overall trend remains apparent even with only 16 taxa (Supplemental Figure 7b).

In addition to examining relative abundances of bacterial taxa between douc groups, we calculated and compared the log *Firmicutes* to *Bacteroidetes* ratio for each douc group (Supplemental Figure 8). The log of the *Firmicutes* to *Bacteroidetes* (F:B) ratio, which has been suggested as a measure of energy harvest capacity by microbial communities (Ley et al. 2006, Turnbaugh et al. 2009, Turnbaugh et al. 2006), was higher in the wild population than in the semi-wild, semi-captive, and captive populations (4.64 ± 0.94; 3.78 ± 1.14; 1.94 ± 0.81; 1.43 ± 0.50, respectively). A Kruskal-Wallis test indicated there is a significant difference in F:B ratio between at least one lifestyle group and the others. Pairwise significance between specific groups, with the exception of captive versus semi-captive (Wilcoxon rank-sum p = 0.05), were significantly different from one another for all lifestyle pairs with p < 0.01. In fact, there appears to be a relationship between lifestyle and the F:B ratio, as we see the highest ratio in wild doucs, the second highest ratio in the semi-wild doucs, the third highest ratio in semi-captive doucs, and finally the lowest ratio in captive doucs (Supplemental Figure 8).

### Red-shanked douc metagenome: Functional analysis using PICRUSt

The functional profiles of the microbial sample in this study were investigated employing PICRUSt. In captivity, we observed a general trend toward increased protein metabolism at the expense of fatty acid metabolism. Specifically, the KEGG Ortholog (KO) super-heading “Amino acid metabolism” was highly correlated with captivity status (polyserial rho = 0.85, p = 2.2x10^-^ 11), and the super-heading “Lipid metabolism” was highly anticorrelated (polyserial rho = −0.89, p = 7.3x10^-12^). Perhaps due to the presence of chloroplasts in the closed-reference data used for PICRUSt analysis, the photosynthesis and antenna proteins pathway was downregulated in captivity, but the porphyrin and chlorophyll metabolism pathway was upregulated. With the exception of tetracycline biosynthesis, antibiotics-related pathways, including vancomycin biosynthesis, beta-lactam resistance, and penicillin & cephalosporin biosynthesis, were positively associated with captivity. Certain xenobiotic (mainly industrial pollutants) degradation pathways were positively associated with captivity, including ethylbenzene, styrene, and toluene. Other xenobiotic pathways (such as plant toxins and wartime chemicals) were negatively associated with captivity, including xylene, dioxins, atrazine, and chloroalkanes & chloroalkenes. Lastly, chemotaxis, invasion, flagellar assembly, and cytoskeleton genes were enriched in wild doucs. All p-values for results in this paragraph were less than < 1x10^-2^ (Figure 5; Supplemental Table 2).

**Figure 5.**
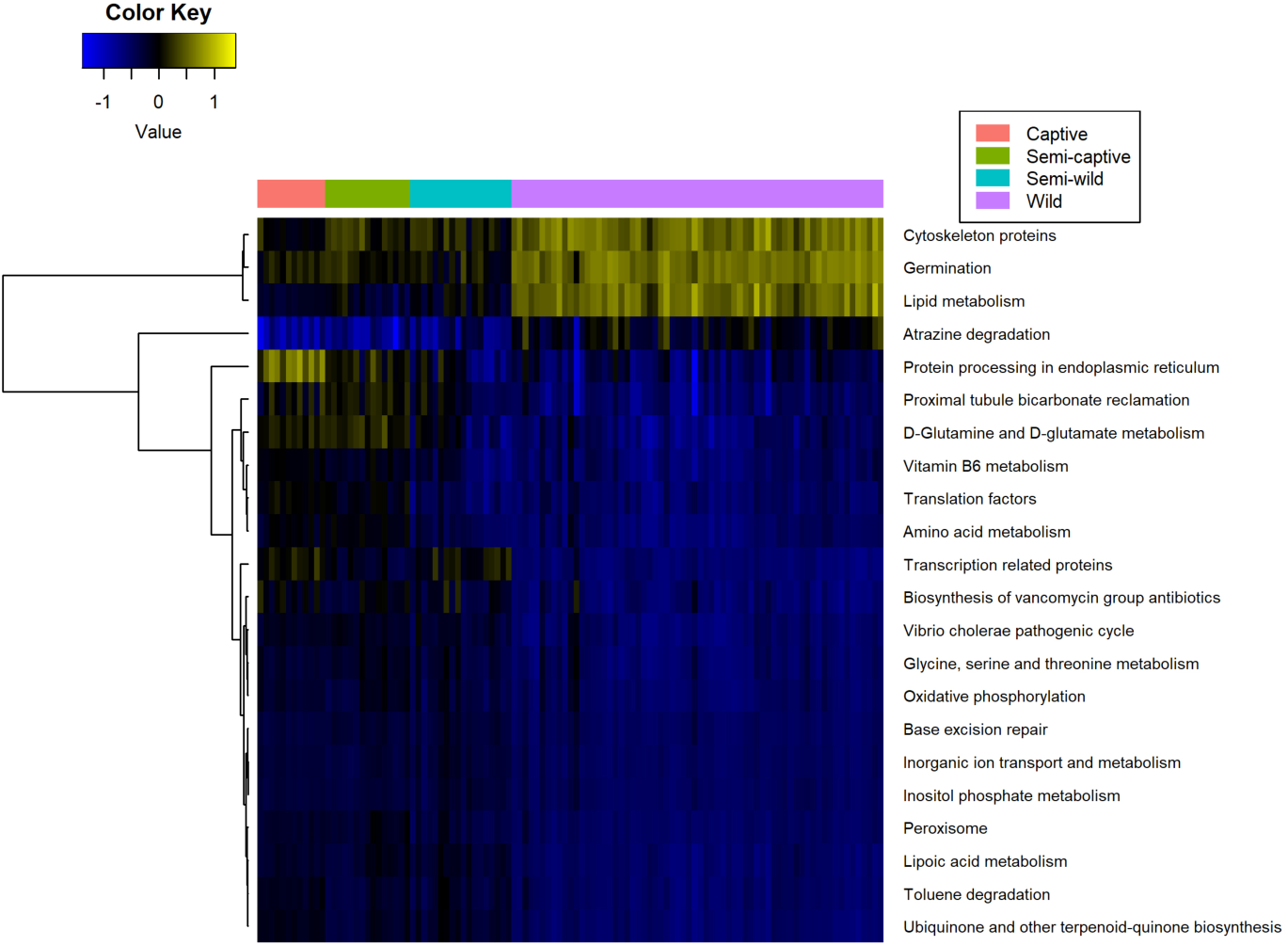
Heatmap of KEGG level 3 metabolic pathways in red-shanked douc groups living four distinct lifestyles. Pathways are displayed with polyserial correlations (rho) above 0.75, rho estimate adjusted p < 0.05, and (pairwise) Wilcoxon rank-sum adjusted p-value for wild and captive lifestyles < 0.05. Color represents intensity of unit-normalized centered log ratio abundances along gradient of color scale shown.

### Composition of douc diets

The diets of the douc populations were compared to determine what factors, if any, could have contributed to the differences in microbiome composition observed (Table 1). Wild doucs fed on 57 different plant species. Sixty-one percent of all identified plant parts observed being ingested were collected and chemically analyzed. The semi-wild douc population were offered 60 plant species over the course of one year, 16 of which were never consumed (Otto 2005). In contrast to the high diet diversity (i.e., number of plant species) consumed by the wild and semi-wild doucs, the semi-captive and captive doucs consumed a much less diverse diet. Specifically, the semi-captive doucs were presented with approximately 15 plant species and the captive douc diet only contained one plant species (Clayton et al. 2016) (Figure 6a). The semi-wild population was observed feeding on 35 different plant genera over one year, and the wild population was observed feeding on 41 different plant genera over approximately seven months. While nine genera were observed being ingested by both the wild and semi-wild populations, only five genera were ingested by both semi-wild and semi-captive doucs. Of the five genera, only one was ingested by wild, semi-wild, and semi-captive doucs. The single plant species consumed by the captive doucs was also consumed by the semi-captive doucs (Figure 6b; Table 2).

**Table 1:** Dietary components of red-shanked doucs living four distinct lifestyles, including wild, semi-wild, semi-captive, and captive.

| Dietary Groups | Proportion of Diet (%) |  |  |  |
| --- | --- | --- | --- | --- |
|  | Wild | Semi-wild | Semi-captive | Captive |
| Leaves* | 65.50 | 100.00 | 66.10 | 1.40 |
| Flowers | 5.30 | 0.00 | 0.00 | 0.00 |
| Seeds | 2.00 | 0.00 | 0.00 | 0.00 |
| Unripe Fruit | 17.80 | 0.00 | 0.00 | 0.00 |
| Other plant parts** | 9.40 | 0.00 | 0.00 | 0.00 |
| Pellets | 0.00 | 0.00 | 0.90 | 7.30 |
| Fruits and Vegetables | 0.00 | 0.00 | 33.00 | 90.90 |
| Cereals | 0.00 | 0.00 | 0.00 | 0.40 |

\* Values for leaves may also include petioles or stems.
\*\*Other plant parts include pith, bark and leaf buds.

**Figure 6.**
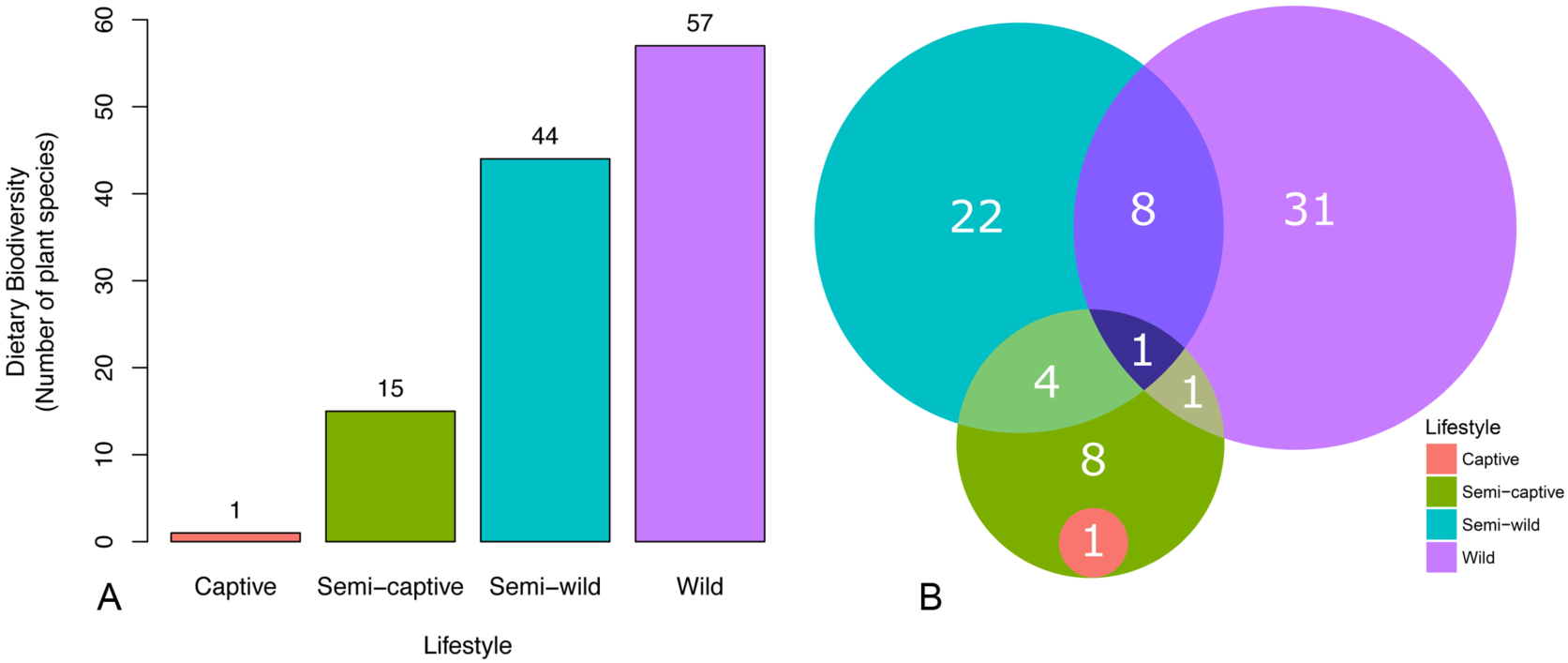
Plant diversity in red-shanked douc diet reflects dietary diversity across populations. (a) Bar plots of dietary biodiversity, as as measured by the number of plant species consumed by wild, semi-wild, semi-captive, and captive populations of red-shanked doucs. Wild doucs feed on 57 different plant species, whereas the semi-wild doucs feed on 44 different plant species annually (Clayton 2015, Otto 2005). In contrast to the high dietary diversity consumed by the wild and semi-wild doucs, semi-captive and captive doucs are fed far fewer plant species. Specifically, semi-captive doucs are feed on approximately 15 plant species and the captive doucs are fed single plant species (Clayton 2015). (b) Venn diagram depicting the number of plant genera consumed by the wild, semi-wild, semi-captive, and captive douc populations, while the numbers in overlaps representing the genera eaten by the constituent populations. Number of genera for the wild population was obtained from Clayton (unpublished). Number of genera for the semi-wild population was obtained from a combination of Clayton (unpublished) and Otto (2005).

**Table 2:** Plant genera observed in douc populations by lifestyle.

| Diets by Lifestyle | Captive | Semi-captive | Semi-wild<sup>1</sup> | Wild |
| --- | --- | --- | --- | --- |
| Plant genus/genera | *\*Morus* | *Acalypha*<br>*\*\*Adenanthera*<br>*Asystasia*<br>*\*\*Cinnamomum*<br>*\*\*Hibiscus*<br>*Khaya*<br>*\*\*Leucaena*<br>*\*\*\*\*\*Litsea*<br>*Moringa*<br>*\*Morus*<br>*Polygonum*<br>*Pterocarpus*<br>*\*\*\*Syzygium*<br>*Tamarindus*<br>*Terminalia* | *\*\*Adenanthera*<br>*Alangium*<br>*Albizzia*<br>*Alchornea*<br>*Averrhoa*<br>*Bidens*<br>*Cassia*<br>*\*\*\*\*Celtis*<br>*\*\*Cinnamomum*<br>*\*\*\*\*Claoxylon*<br>*Clerodendrum*<br>*Crateva*<br>*\*\*\*\*Dalbergia*<br>*Delonix*<br>*Derris*<br>*Dimocarpus*<br>*Elaeocarpus*<br>*Euodia*<br>*\*\*\*\*Ficus*<br>*\*\*Hibiscus*<br>*\*\*\*\*Ilex*<br>*\*\*Leucoena*<br>*\*\*\*\*\*Litsea*<br>*Maesa*<br>*Micromelum*<br>*Mussaenda*<br>*\*\*\*\*Oroxylon*<br>*Phyllanthus*<br>*Rhus*<br>*Saurauia*<br>*Sterculia*<br>*Triadica*<br>*\*\*\*\*Vitex*<br>*Wrightia*<br>*\*\*\*\*Zanthoxylum* | *Acacia*<br>*Amesiodendron*<br>*Ancistrocladus*<br>*Antidesma*<br>*Aporosa*<br>*Barringtonia*<br>*Beilschmiedia*<br>*Bischofia*<br>*Brownlowia*<br>*Callerya*<br>*\*\*\*\*Celtis*<br>*\*\*\*\*Claoxylon*<br>*Cleidion*<br>*\*\*\*\*Dalbergia*<br>*Diospyros*<br>*Dipterocarpus*<br>*\*\*\*\*Ficus*<br>*Glochidion*<br>*Gluta*<br>*Gmelina*<br>*\*\*\*\*Ilex*<br>*Litchi*<br>*Lithocarpus*<br>*\*\*\*\*\*Litsea*<br>*Mallotus*<br>*Millettia*<br>*Mischocarpus*<br>*Nauclea*<br>*Ochrocarpus*<br>*Ormosia*<br>*\*\*\*\*Oroxylon*<br>*Parashorea*<br>*Quercus*<br>*Rothmannia*<br>*Sandoricum*<br>*Schefflera*<br>*Scolopia* |
|  |  |  |  | *** *Syzygium*<br>*Uvaria*<br>**** *Vitex*<br>**** *Zanthoxylum* |

<sup>1</sup>Combination of data collected by JBC for this study and data lifted from Otto (2005)
\*Shared between captive and semi-captive
\*\*Shared between semi-captive and semi-wild
\*\*\*Shared between semi-captive and wild
\*\*\*\*Shared between semi-wild and wild
\*\*\*\*\*Shared between semi-captive, semi-wild, and wild

We estimated total raw plant dietary content for the four douc populations, using chloroplast sequences observed in the 16S amplicon sequencing data. We found that chloroplast content was substantial in wild and semi-wild populations, but that chloroplast content decreased dramatically in semi-captive and captive doucs (Figure 7a). Alignment of the 16S sequences to known plant reference genomes at 95% identity yielded a heatmap similar to Supplemental Figure 5b. Overall, there is a clear trend toward increased plant abundances in the semi-wild and wild lifestyles, although certain orders display different abundances between lifestyle groups. Some orders can be seen to overlap between lifestyles, while others do not (Figure 7b; Supplemental Table 3).

**Figure 7.**
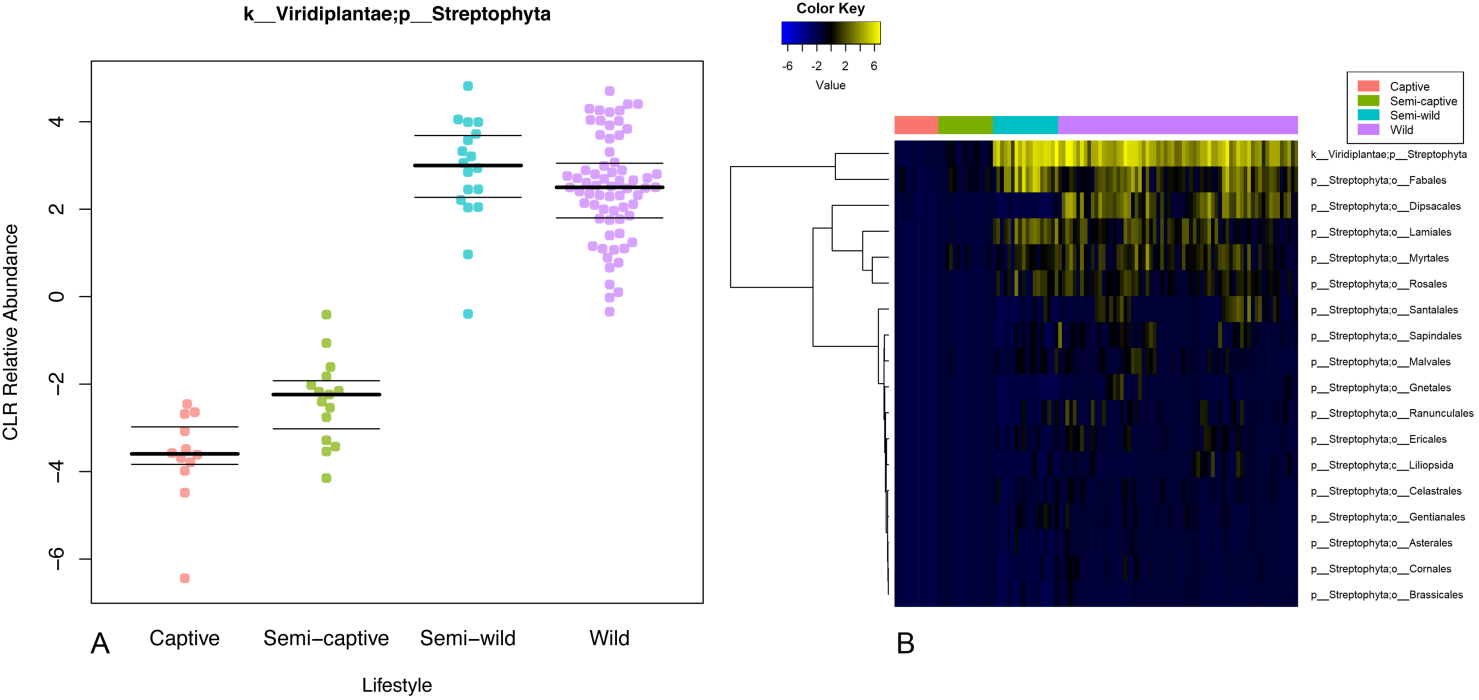
Plant chloroplast material in douc feces increases with wildness of lifestyle. (A) Centered-log-ratio-corrected relative abundance beeswarm plot. (B) Heatmap of class/order-level representation of plant taxa by lifestyle. All land plants (phylum Streptophyta) for which refseq chloroplast sequences were available were searched for matches with the 16S data. There is trend toward increased plant matter with wildness (polyserial rho = −0.81, pairwise wild-captive Wilcoxon rank-sum adjusted p = 8.65×10^−13^). Certain plant taxa display different patterns of overlapping abundance between lifestyles.

Nutritional composition of food items included in wild, semi-wild, semi-captive, and captive douc diets differed. Specifically, the crude protein concentration of the semi-wild, semi-captive, and captive douc diets was higher than that of the wild douc diet. The wild and semi-wild douc diets contained much more Acid Detergent Fiber (ADF) and Neutral Detergent Fiber (NDF) than did the semi-captive and captive diets. We examined the four douc diets for differences in amount of three macrominerals, including calcium, potassium, and sodium. Of the diets examined, the semi-wild douc diet contained more calcium than did the wild, semi-captive, and captive douc diets. Additionally, the diet consumed by wild doucs contained more potassium than did the diets consumed by semi-wild, semi-captive, or captive doucs. The diets of semi-wild and wild doucs contained considerably less sodium than the semi-captive and captive doucs. The semi-wild douc diet contained more iron and zinc than the wild, semi-captive, or captive diets. The concentration of sugar was not available for all lifestyle groups. The captive douc diet had the highest amount of soluble sugars compared to wild and semi-wild diets (Table 3).

**Table 3:**
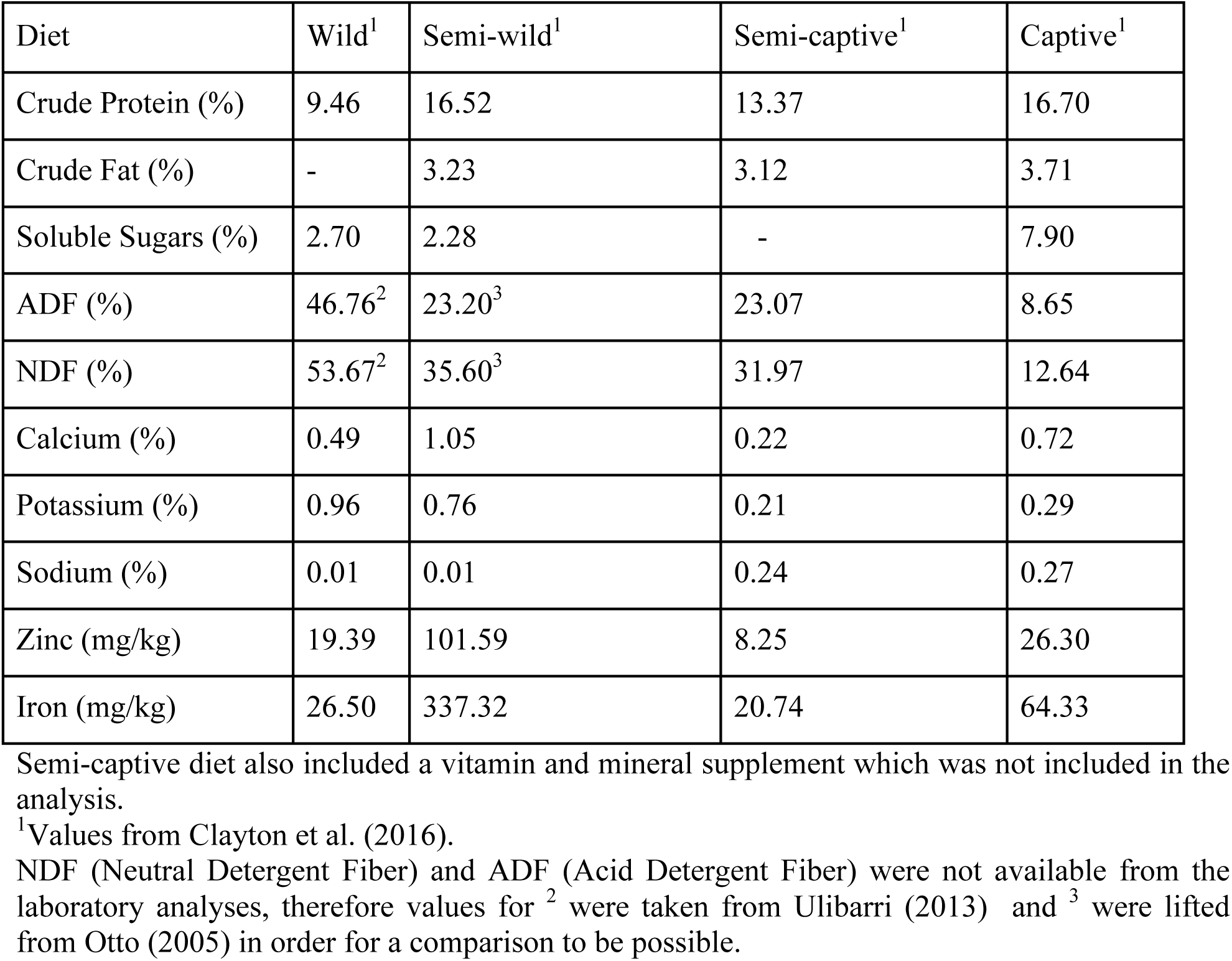
Nutrient content on a dry matter basis from red-shanked doucs living four distinct lifestyles, wild, semi-wild, semi-captive, and captive.

## DISCUSSION

In this study, the red-shanked douc was used as a model system to study the relationships between dietary composition and microbial composition and function within the gastrointestinal tract. Doucs are folivorous Old World monkeys, that are anatomically, physiologically, and ecologically unique amongst the living primates (Davies and Oates 1994). They possess specialized GI systems similar to ruminants, allowing for the digestion and utilization of extremely high fiber diets (Chivers 1994, Lambert 1998). For doucs, mutualistic microbial populations are indispensable to digestive processes such as the fermentation of polysaccharides and subsequent production of short-chain fatty acids (Jablonski 1998, Nijboer et al. 2006, Nijboer, Clauss, and Olsthoorn 2006). Although the digestive specializations possessed by doucs have allowed them to thrive in their native habitat, the same specializations appear to challenge their survival in captivity, as they are highly susceptible to gastrointestinal disorders when maintained on commercially prepared diets in captive situations (Agoramoorthy, Alagappasamy, and Hsu 2004, Nijboer 2006, Power, Toddes, and Koutsos 2012). We hypothesized that specific and unique microbial subsets play a critical role in the utilization of fibrous vegetation with natural toxicants, and that captive doucs lack the microbiota to maintain optimal health due to inadequate dietary substrate. In order to better understand the link between lifestyle, gut microbial communities, and health, we examined the fecal microbiomes of four douc populations living four distinct lifestyles (wild, semi-wild, semi-captive, and captive).

### Microbial diversity

Studies have shown that present-day (i.e., modern) humans have lost a considerable portion of their natural (i.e., historical) microbial diversity (Clemente et al. 2015, Martinez et al. 2015, Moeller et al. 2014). Reduced bacterial diversity is often viewed as a negative indicator of health (Fujimura et al. 2010). 16S rRNA sequencing results revealed that captive doucs had a marked reduction in gut bacterial alpha diversity when compared to wild and semi-wild doucs, as we reported recently (Clayton et al. 2016). Considering that doucs often fail to thrive in captivity, this was a salient finding. Not only was a reduction in diversity detected in captive doucs, but a gradient-like decrease in diversity related to lifestyle was observed, as the level of diversity observed in the semi-wild doucs was intermediate between wild and semi-captive doucs, and the level of diversity seen in semi-captive doucs was intermediate between semi-captive and captive doucs. This trend is consistent between metrics of richness and evenness, and suggest that lifestyle factors, especially dietary composition, and gut bacterial diversity are interrelated for doucs. This is similar to what has been shown in humans and other organisms (Clemente et al. 2015).

Beta-diversity metrics revealed each douc population had a unique microbiome. Weighted UniFrac ordination, although maintaining clear group separation, did not recover a clear gradient, whereas Bray-Curtis appears similar to Unweighted UniFrac in visibly resolving the lifestyle gradient with PC1. This implies that the taxonomic membership of the gut microbiome, more than the abundance of each member, plays a dominant role in uncovering the gradient between lifestyles. Moreover, tree-independent unsupervised hierarchical clustering, utilizing only the taxonomic features significantly correlated to lifestyle gradient, preserved lifestyle group membership without erroneously assigning any samples to the wrong lifestyle groups, and found further structure within each group. This highlights the potential for using these taxa, or a subset thereof, as biomarkers capable of accurately differentiating between lifestyles.

### Lifestyle and diet drive gut microbial community structure

Establishing a link between diet and the microbiome was a major focal point of this study. GI microbiome composition is shaped by host genetics and environment, among many factors (David et al. 2014, Goodrich et al. 2014). Examining four populations of the same NHP species living in four very different environments enabled assessment of the contribution of environmental factors independent of interspecific host variation towards shaping the microbiome. The major environmental differences to which each douc population is exposed, such as climate and diet, suggest that environment plays a fundamental role in shaping gut microbiome composition in wild and captive NHPs. Of the environmental factors that contribute to the establishment and maintenance of the gut microbiota, diet is likely the most influential, as studies have shown that changes in diet are directly associated with shifts in gut microbial community structure (David et al. 2014, Gophna 2011, Muegge et al. 2011, Wu et al. 2011, Xu and Knight 2015). Examples exist of species adapting to specific dietary niches in both wild (Amato et al. 2014) and captive (Kohl, Skopec, and Dearing 2014) settings via changes in their gut microbiota.

Many of our results suggest the existence of a relationship between microbiome composition and dietary patterns. The relative abundance of select bacterial genera, *Bacteroides*, *Prevotella*, *Oscillospira*, and *Blautia*, and differences in dietary composition between lifestyles, warrant specific discussion in this regard. *Prevotella*, which is involved in the digestion of simple sugars and carbohydrates (Purushe et al. 2010), was notably higher in the captive doucs than in the other three douc populations examined. One very different component in the diets of wild versus captive doucs is the inclusion of produce in the captive diets. Fruits consumed by captive primates have much different nutrient profiles than fruits consumed by wild primates (Oftedal and Allen 1996). They have been cultivated for human consumption to be lower in fiber and protein and higher in sugar as opposed to wild fruits which are, in general, the exact opposite (Schwitzer and Kaumanns 2003). Given the diet of captive doucs contained more than a threefold increase in the percentage of sugars compared to wild and semi-wild douc diets, there seems to be a clear relationship between sugar consumption and *Prevotella* abundance. Unlike captive doucs, wild doucs consumed unripe fruit, which is drastically different than ripe fruit fed to captive individuals, and thus its impact on the douc microbiome is different than cultivated fruits would have. The low relative abundance of *Prevotella* in the wild douc microbiome provides further evidence that diet is a major driver of microbiome composition.

Semi-captive and captive doucs also harbored more *Bacteroides* than semi-wild or wild doucs. In humans, *Bacteroides* are found in higher abundance in individuals who consume diets high in fat and protein (Wu et al. 2011, Yatsunenko et al. 2012). Interestingly, the captive douc diet contained more protein than the wild douc diet, which may explain why *Bacteroides* was a dominant member of the captive douc microbiome, yet virtually absent from the wild douc microbiome.

Another notable example of this diet-microbiome relationship seen in our analysis was with the genus *Oscillospira*, which has a known association with the consumption of plant material, including leaves and grass cuticles (Clarke 1979, Mackie et al. 2003, Yanagita et al. 2003, Zoetendal et al. 2013). *Oscillospira* was markedly increased in the wild doucs compared to the other douc populations, and was more abundant in the semi-wild and semi-captive doucs than in the captive population. The observed differences in abundance of *Oscillospira* between douc populations was likely a function of the stark differences in dietary consumption between populations, and most importantly, the difference in diversity and proportion of plants and plant parts consumed by the different douc populations examined. Wild, semi-wild, and, to a lesser degree, semi-captive doucs all consume diets that contain a higher proportion and diversity of plants compared to the captive population. Aside from *Oscillospira*, overall douc microbiome composition seemed to be, at the very least, partially driven by plant abundance and diversity in the diet. While a high variety of plant species is bound to impact the gut microflora, the sheer differences in proportion of the diet which is plants is likely to be equally as much of a causative factor (de Menezes et al. 2011). In this study, we utilized both known dietary makeup and measured chloroplast content (via 16S rRNA sequencing) to detail plant consumption by each douc population. We observed a striking downward trend in the number of plant genera and species consumed by increasing captivity of lifestyle (Figure 6a; Figure 6b). The measured chloroplast content of the stool mirrored this downward trend (Figure 7a). Further, we were able to observe trends in plant taxon composition, although the resolution of the chloroplast analysis was limited approximately to the plant class level. Some of the patterns of overlap observed mirrored the overlap in plant genera fed to the doucs, but we were unable to confirm whether the presence of the plant classes/orders reported in the chloroplast analysis indeed coincided with the specific plants fed to the doucs in the different lifestyles. Further targeted plant genomic screens may expand upon this proof of concept in the future.

An unexpected result found in this study was the high abundance of the genus *Akkermansia* found in the semi-wild doucs. While *Akkermansia* was most abundant in the semi-wild doucs by far, this genus was also more abundant in the wild doucs than doucs living semi-captive and captive lifestyles. Members of the genus *Akkermansia*, such as *Akkermansia muciniphila*, are known for their roles in mucin-degradation, and have been suggested to play protective roles in the gut (Belzer and de Vos 2012, Everard et al. 2013). Everard et al. (2013) showed that obese and type 2 diabetic mice had decreased abundance of *A. muciniphila*, and treatment with this microbe reversed high-fat diet-induced metabolic disorders. Interestingly, a recent study examining the link between gut microbiota and primate GI health found GI-unhealthy doucs had reduced relative abundances of *Akkermansia* (Amato et al. 2016). Another study examining gut microbiome composition of a colobine primate, *Rhinopithecus brelichi*, showed that *Akkermansia* was more abundant in captive individuals when compared to their wild counterparts, which is different than what was seen in doucs (Hale 2014). The extremely high abundance of *Akkermansia* in the semi-wild doucs combined with the higher level seen in wild doucs compared to semi-captive and captive doucs suggests that the microbe is linked to diet, as the diets of wild and semi-wild doucs contain much more plant diversity compared to those of the captive doucs.

We examined the F:B ratio, as this ratio is important in humans in terms of dietary energy extraction (Ley et al. 2008, Turnbaugh et al. 2006). We saw a higher F:B ratio in wild and semi-wild doucs compared to semi-captive and captive doucs. Ley et al. (2006) found an increased presence of Firmicutes with a corresponding decrease of Bacteroidetes correlating with an overall greater energy harvest (Ley et al. 2006). Based on our results, it appears a decrease in the F:B ratio was clearly associated with lifestyle, notably diet, as the wild doucs had the highest ratio, followed by the semi-wild doucs, semi-captive doucs, and captive doucs. As previously mentioned, wild and semi-wild diets contained substantially more plant matter than captive diets. Naturally this equates to diets much higher in fiber fractions (ADF, NDF). Due to the scarcity of high-quality food items in the wild, we witnessed doucs ingesting very fibrous plant parts such as bark, mature leaves, flowers, seeds and unripe fruit. In the semi-wild facility, the doucs are habituated and know that they will receive leaf meals which provides them with a balance of fiber and soluble nutrients, making the ingestion of very fibrous items such as bark unnecessary. This can partially explain the higher reported NDF values in wild doucs when compared to semi-wild doucs. This relationship between F:B ratio and diet was expected, as our results show captive populations have diets lower in fiber fractions and higher in soluble carbohydrates, notably sugars, when compared to wild or semi-wild populations. Overall, the differences in the F:B ratio observed between populations living in natural versus unnatural settings, suggests the ratio is an indicator of overall gut health, as a higher ratio is associated with a higher fermentation efficiency and increased VFA production (Amato et al. 2014, Turnbaugh et al. 2006), and doucs living under artificial (i.e., captive) conditions, which had a lower ratio, often suffer from a wasting syndrome (Crissey and Pribyl 1997, Lacasse et al. 2007).

### Putative functional associations with lifestyle

PICRUSt-predicted functional pathways show a few interesting trends. Interestingly, captivity appears to be correlated with pathways spanning metabolism of antibiotics, nutrients, and xenobiotics, as well as other potentially relevant trends. In terms of antibiotics, beta-lactam antibiotic resistance genes were enriched in captivity alongside its production (penicillin and cephalosporin biosynthesis) in captive lifestyles. Tetracycline resistance genes, in contrast, were elevated in the wild without an attendant significant upregulation of resistance. This pattern raises the possibility that the antibiotics pathways upregulated in captivity may be adapted for competition and virulence factor regulation (Balasubramanian et al. 2011), whereas the pathways upregulated in the wild may play more of a quorum sensing role (Lu 2006).

Another interesting trend is the apparent tradeoff between amino acid metabolism in the wild and lipid metabolism in captivity, two important umbrella pathways (hierarchically from top level of KEGG, Metabolism -> Lipid Metabolism and Metabolism -> Amino Acid Metabolism). This pronounced trend is likely indicative of the highly different diets received by the populations and may reflect differences in plant species consumed, the differences in microbiome composition in response to different nutrient profiles, or other lifestyle factors including antibiotics exposure. Additionally, the differentially increased motility of the members of the wild microbiome (evidenced by upregulated cytoskeletal regulation, flagellar and motility proteins, and chemotaxis) may imply increased vigor and facilitate nutrient scavenging. The increasing differential abundance of sporulation and germination pathways with wildness may also highlight the resilience of the wild microbiota, and may be due in part to various members of order Clostridiales (many of which can form spores and later germinate from them) also being differentially more abundant.

There are some particularly intriguing trends concerning xenobiotic (pollutant) degradation. Xylene, dioxins, atrazine, and chloroalkane/ene degradation levels are significantly associated with increasing wildness of lifestyle. Setting aside the expected presence of toxic chemicals in the douc’s natural diet, all of these compounds have been associated with wartime chemicals. Specifically, agent orange and other war chemicals deployed during the Vietnam Conflict are atrazines and dioxins. Dioxins are also byproducts of forest fires and several types of manufacturing processes. They are persistent in the environment (Schecter et al. 2006), and studies have shown that manufacturing workers who are in direct contact with dioxins have an increased risk for the development of cancer (Kogevinas et al. 1995). Son Tra Nature Reserve, the research site where wild douc fecal samples were collected, is located approximately 8 km from Danang International Airport. This airport is considered one of the world’s most dioxin-contaminated sites. There are over 187,000 square meters of contaminated soil located in several sites near the airport (Minh et al. 2009). Similarly, jet fuel and industrial solvents often contain xylene, another xenobiotic upregulated in wilder lifestyles. The semi-wild population, which shows a peak in xylene degradation, is located in Singapore, which contains the largest xylene plant (Tremblay 2011). When refined into xylyl, xylene was also used as a wartime riot control agent (Olajos and Stopford 2004). Interestingly, the reverse (i.e. showing an increase with captivity) is observed for the degradation pathways of other pollutants often associated with industrialized economies such as ethylbenzenes (Welch and Fallon 2005), styrenes (often industrially synthesized from ethylbenzenes) (James and Castor 1994), and toluene (Fishbein 1987), which may indicate forms of air contamination or other “first-world” pollutants. Given this, coupled with the fact that wild doucs consume plants rich in toxic compounds, we hypothesize that the microbiome of wild red-shanked doucs serves a detoxification role for the animal and therefore is enriched for taxa with the ability to degrade local contaminants. Another interesting implication is that the microbiome may function as a geochemical sensor of environment, where passage through a host may modulate detection sensitivity for certain compounds through dietary biomagnification.

### Viewing the microbiome as compositional data

From a statistical standpoint, the treatment of microbiome data in a modern compositional framework (Gloor et al. 2016) frees microbial composition data from the simplex and allows many standard univariate statistical analysis techniques to apply (Van den Boogaart and Tolosana-Delgado 2013). With the observation that most bacterial abundances post-CLR transformation appeared to be roughly gaussian, the polyserial correlation test became applicable to assess the degree by which each microbial or functional pathway abundance was correlated with the latent continuous variable captured by the four ordered “lifestyle” categories. For additional conservatism, the nonparametric Wilcoxon rank-sum test was also used on the two extrema of the assumed gradient (the “Wild” and “Captive” lifestyles), and the significance criterion for an association was amended to require both strong polyserial correlation (absolute rho above 0.3) as well as corrected Wilcoxon p-values less than 0.05. Importantly, this compositional framework allows for the highly-powered and gradient-centric interpretation and visualization of microbial and functional data.

By modeling the lifestyle groups themselves as a latent gradient variable, we are assessing microbial and functional relationships with a composite metric of “wildness” or “lifestyle perturbation,” and hence avoid the use of other potentially confounded covariates such as climatic factors, diversity of plant species consumed, frequency of human contact, or health markers as individual proxies. In so doing, these and other covariates can be interpreted in relation to one another. Our focus in this work was to ascertain which microbes and microbially-deduced functional pathways were correlated with lifestyle and interpret these correlations in context of collected health and dietary data.

This analysis revealed a selection of strongly correlated, statistically significant microbial biomarkers indicative of the health and well-being of the red-shanked douc across lifestyle conditions. These trends may have implications to human and livestock health, as the douc may serve as a genetically similar model for the former, and a digestively similar model for the latter. Since the microbiome itself is associated with host genetics as well as digestive functions and diseases (Knights et al. 2014), this study provides the framework for and invites further investigation of this potential model organism for the applicability of these findings within other species.

## ACKNOWLEDGEMENTS

We thank the Endangered Primate Rescue Center, Singapore Zoo, and Philadelphia Zoo for providing fecal samples from semi-wild, semi-captive, and captive red-shanked doucs; Tran Van Luong, Nguyen Van Bay, and Nguyen Manh Tien for their permission to work in Son Tra Nature Reserve and for their continued support and help; Kieu Thi Kinh and Thai Van Quang for help in obtaining the research permits; the Department of Forest Protection, the Danang University, and the Son Tra Nature Reserve for granting the research permits; and Christina Valeri and James Collins at the University of Minnesota Veterinary Diagnostic Laboratory for their assistance with acquiring and maintaining shipping permits. This research was funded in part by the Margot Marsh Biodiversity Foundation; the Mohamed bin Zayed Species Conservation Fund; and the National Institutes of Health through a PharmacoNeuroImmunology Fellowship (NIH/National Institute on Drug Abuse T32 DA007097-32) awarded to JBC.

## SUPPLEMENTAL TABLES

**Supplemental Table 1:**
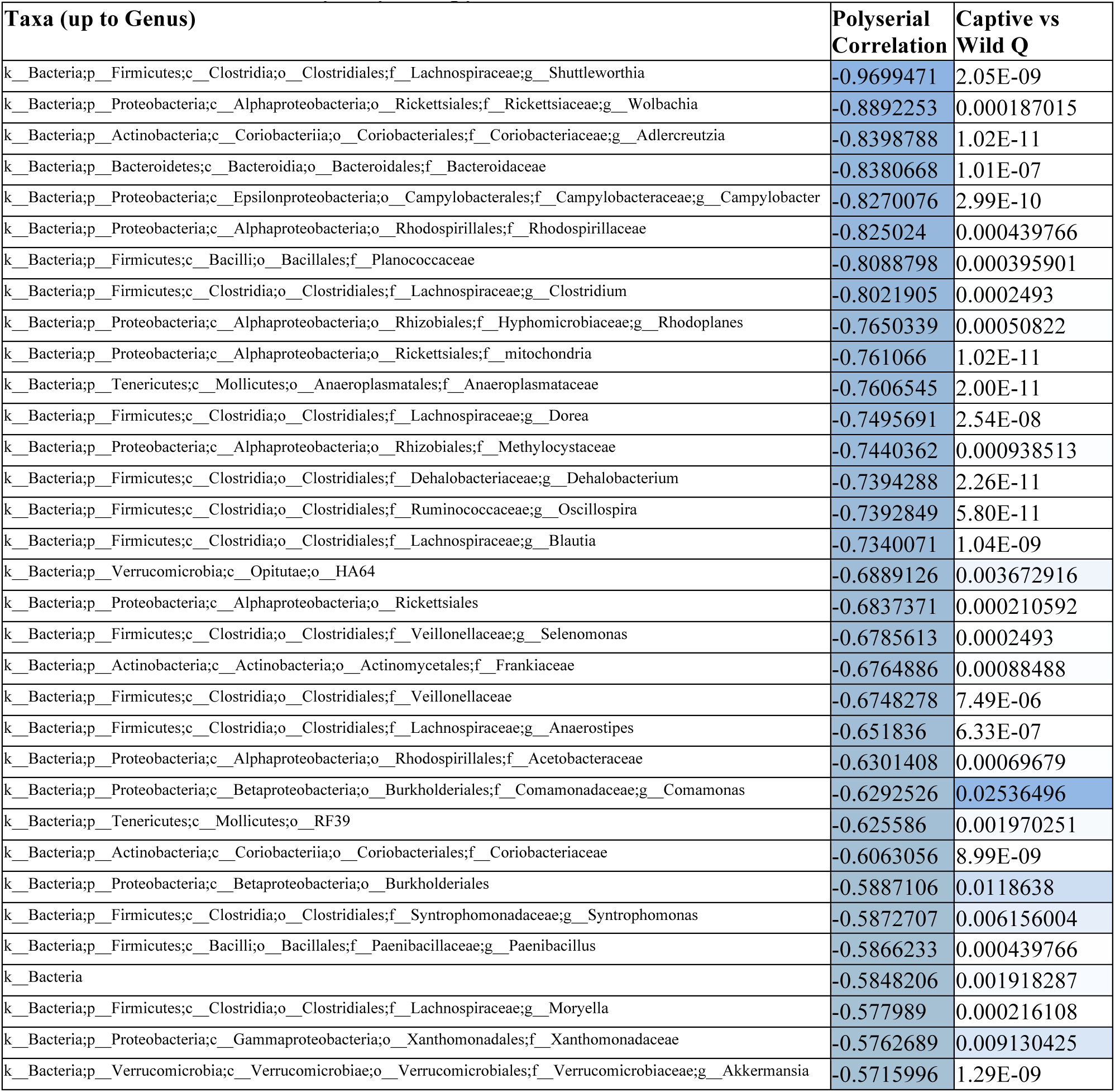
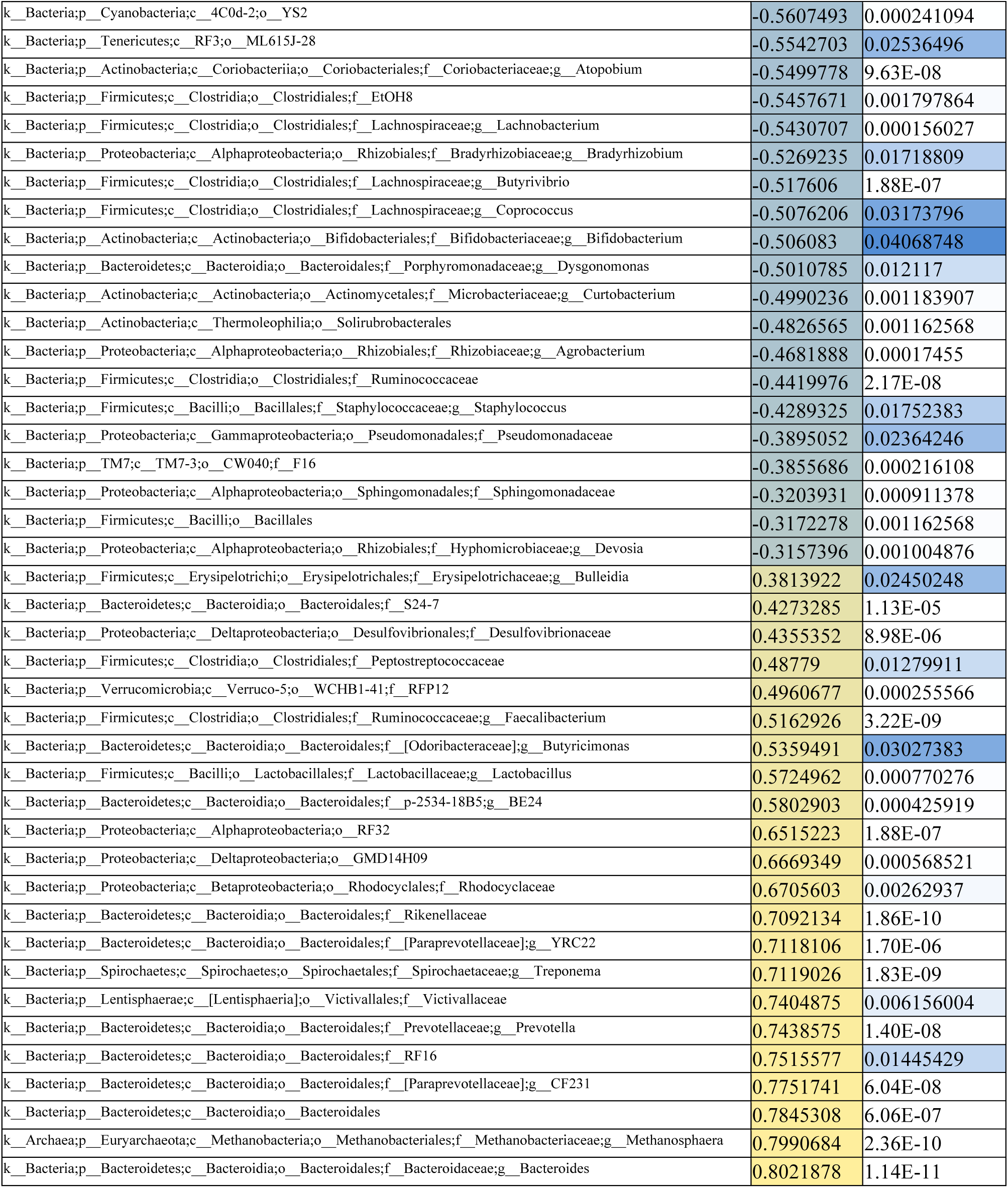
Differentially abundant genera. Statistical significance of differentiation was assessed using pairwise Wilcoxon rank-sum tests of each taxon’s centered-log-ratio transformed abundance in the Captive and Wild lifestyles, as well as the polyserial correlation of the same across all four lifestyles as the ordered factor Wild < Semi-wild < Semi-captive < Captive. The polyserial correlation column is colored according to intensity of correlation; blue signifies decreased abundance with captivity level and yellow increased abundance with captivity level. For the adjusted p-values (Q value column), intensity of color corresponds to Wilcoxon p-value up to alpha = 0.05. Criteria for display included having an adjusted Wilcoxon rank-sum p < 0.05, absolute polyserial correlation above 0.3, and polyserial rho p-value < 0.05. Taxa are displayed at the most specific taxonomic level to which they were annotated; taxa lacking annotation at the genus level can be interpreted as being “other” genera within the level of taxonomy they occupy.

**Supplemental Table 2.** Differentially abundant functional pathways. Statistical significance of differentiation was assessed using pairwise Wilcoxon rank-sum tests of each pathway’s centered-log-ratio transformed abundance in just the Captive and Wild lifestyles, as well as the polyserial correlation of the same across all four lifestyles as the ordered factor Wild < Semi-wild < Semi-captive < Captive. The polyserial correlation column is colored according to intensity of correlation; blue signifies decreased abundance with captivity level and yellow increased abundance with captivity level. For the adjusted p-values (Q value column), intensity of color corresponds to Wilcoxon p-value up to alpha = 0.05. Criteria for display included having an adjusted Wilcoxon rank-sum p < 0.05, absolute polyserial correlation above 0.3, and polyserial rho p-value < 0.05.

| Pathway | Polyserial Correlation | Captive vs Wild Q |
| --- | --- | --- |
| Ubiquinone and other terpenoid-quinone biosynthesis | 0.86613 | 1.65E-11 |
| Vibrio cholerae pathogenic cycle | 0.8522796 | 1.76E-10 |
| Amino acid metabolism | 0.8485574 | 2.21E-11 |
| Toluene degradation | 0.8355032 | 2.63E-11 |
| Lipoic acid metabolism | 0.8349192 | 1.47E-11 |
| Proximal tubule bicarbonate reclamation | 0.8285716 | 2.34E-11 |
| Peroxisome | 0.8264319 | 2.34E-11 |
| D-Glutamine and D-glutamate metabolism | 0.809642 | 1.32E-11 |
| Transcription related proteins | 0.8057511 | 1.95E-11 |
| Translation factors | 0.8016383 | 1.40E-11 |
| Protein processing in endoplasmic reticulum | 0.7868988 | 2.62E-10 |
| Inositol phosphate metabolism | 0.7778543 | 1.86E-10 |
| Vitamin B6 metabolism | 0.7715443 | 2.96E-11 |
| Base excision repair | 0.7714182 | 2.92E-10 |
| Biosynthesis of vancomycin group antibiotics | 0.7689167 | 2.76E-10 |
| Inorganic ion transport and metabolism | 0.7637347 | 1.60E-10 |
| Glycine, serine and threonine metabolism | 0.7581197 | 7.85E-10 |
| Oxidative phosphorylation | 0.7568452 | 6.65E-10 |
| beta-Lactam resistance | 0.7432696 | 1.91E-09 |
| Phosphatidylinositol signaling system | 0.739012 | 6.11E-10 |
| Isoflavonoid biosynthesis | 0.7284541 | 2.31E-10 |
| Alanine, aspartate and glutamate metabolism | 0.7259672 | 3.26E-09 |
| Tyrosine metabolism | 0.7253912 | 2.92E-10 |
| Purine metabolism | 0.7253711 | 8.19E-09 |
| Metabolism of cofactors and vitamins | 0.7236583 | 6.63E-09 |
| Function unknown | 0.7168606 | 6.63E-09 |
| Ascorbate and aldarate metabolism | 0.7161581 | 5.06E-09 |
| Protein folding and associated processing | 0.7070021 | 3.26E-09 |
| Glutamatergic synapse | 0.7026025 | 4.82E-11 |
| Ubiquitin system | 0.7021036 | 9.09E-11 |
| One carbon pool by folate | 0.694698 | 3.15E-11 |
| General function prediction only | 0.6884648 | 2.14E-08 |
| Polyketide sugar unit biosynthesis | 0.685304 | 1.32E-08 |
| Nucleotide excision repair | 0.6826238 | 1.92E-08 |
| Other ion-coupled transporters | 0.677045 | 8.62E-09 |
| Phosphonate and phosphinate metabolism | 0.6754849 | 1.86E-10 |
| Sphingolipid metabolism | 0.6597281 | 6.30E-09 |
| Valine, leucine and isoleucine degradation | 0.6583393 | 1.07E-10 |
| Aminobenzoate degradation | 0.6487151 | 1.68E-10 |
| Renal cell carcinoma | 0.6353665 | 3.17E-08 |
| Styrene degradation | 0.6319073 | 5.33E-09 |
| Retinol metabolism | 0.6318299 | 1.02E-07 |
| Lysine degradation | 0.6300815 | 1.68E-10 |
| Ethylbenzene degradation | 0.627926 | 4.13E-07 |
| beta-Alanine metabolism | 0.6127058 | 1.84E-08 |
| Huntington's disease | 0.6073938 | 7.66E-10 |
| Primary immunodeficiency | 0.5987362 | 2.05E-06 |
| Bacterial secretion system | 0.5980548 | 1.57E-06 |
| Tryptophan metabolism | 0.5884344 | 4.71E-10 |
| Carbohydrate digestion and absorption | 0.5876023 | 5.04E-05 |
| Glyoxylate and dicarboxylate metabolism | 0.58556 | 1.76E-07 |
| Geraniol degradation | 0.5758705 | 3.44E-08 |
| Pentose and glucuronate interconversions | 0.574738 | 2.94E-06 |
| Proteasome | 0.5693384 | 1.14E-09 |
| Glutathione metabolism | 0.5680232 | 2.93E-09 |
| Primary bile acid biosynthesis | 0.5644047 | 1.90E-05 |
| Bacterial toxins | 0.5640835 | 3.57E-07 |
| DNA repair and recombination proteins | 0.5619595 | 6.34E-07 |
| Butanoate metabolism | 0.5611606 | 1.43E-05 |
| Sulfur metabolism | 0.5512656 | 9.22E-06 |
| Carbon fixation in photosynthetic organisms | 0.5487902 | 7.47E-05 |
| Cell cycle - Caulobacter | 0.5453256 | 6.84E-06 |
| Influenza A | 0.5451721 | 3.07E-09 |
| Arginine and proline metabolism | 0.5397563 | 4.52E-07 |
| Renin-angiotensin system | 0.5395988 | 8.61E-05 |
| Peptidases | 0.5366534 | 2.69E-06 |
| Secondary bile acid biosynthesis | 0.5348466 | 0.000119449 |
| Pathways in cancer | 0.5310567 | 1.59E-07 |
| Porphyrin and chlorophyll metabolism | 0.5304321 | 2.36E-06 |
| Penicillin and cephalosporin biosynthesis | 0.5278086 | 1.32E-07 |
| Epithelial cell signaling in Helicobacter pylori infection | 0.5246982 | 2.58E-08 |
| Taurine and hypotaurine metabolism | 0.5205521 | 2.64E-06 |
| Fatty acid metabolism | 0.5196525 | 7.33E-07 |
| Linoleic acid metabolism | 0.514908 | 2.21E-05 |
| Limonene and pinene degradation | 0.5124961 | 2.25E-07 |
| Pyruvate metabolism | 0.5080889 | 8.66E-06 |
| Betalain biosynthesis | 0.5051249 | 1.12E-07 |
| Caprolactam degradation | 0.5033751 | 1.84E-08 |
| Bladder cancer | 0.5029604 | 2.16E-07 |
| Arachidonic acid metabolism | 0.4984154 | 0.03065246 |
| Indole alkaloid biosynthesis | 0.4961972 | 8.62E-09 |
| Chlorocyclohexane and chlorobenzene degradation | 0.4911376 | 1.48E-07 |
| Adipocytokine signaling pathway | 0.4900165 | 4.52E-07 |
| Meiosis - yeast | 0.4840796 | 4.69E-07 |
| Peptidoglycan biosynthesis | 0.4716894 | 8.99E-08 |
| Hypertrophic cardiomyopathy (HCM) | 0.4646138 | 4.78E-06 |
| Glycosphingolipid biosynthesis - lacto and neolacto series | 0.463966 | 6.35E-06 |
| Bisphenol degradation | 0.4605462 | 0.000143559 |
| Apoptosis | 0.4585675 | 6.01E-08 |
| Amyotrophic lateral sclerosis (ALS) | 0.4543189 | 8.12E-06 |
| Metabolism of xenobiotics by cytochrome P450 | 0.4531665 | 1.56E-05 |
| Drug metabolism - cytochrome P450 | 0.4502912 | 9.81E-06 |
| Circadian rhythm - plant | 0.4451969 | 0.000116798 |
| Naphthalene degradation | 0.4430097 | 0.002843726 |
| Amino acid related enzymes | 0.4348306 | 0.000212966 |
| Protein export | 0.4147701 | 0.000265293 |
| Propanoate metabolism | 0.4059516 | 0.002464333 |
| Amino sugar and nucleotide sugar metabolism | 0.3813197 | 0.004732689 |
| Selenocompound metabolism | 0.3793765 | 0.007380699 |
| Tuberculosis | 0.3765017 | 0.000983163 |
| Stilbenoid, diarylheptanoid and gingerol biosynthesis | 0.3715535 | 0.00017531 |
| Biosynthesis of unsaturated fatty acids | 0.364953 | 0.006819423 |
| Fluorobenzoate degradation | 0.3606798 | 0.003021935 |
| Synthesis and degradation of ketone bodies | 0.3599332 | 0.008969831 |
| Galactose metabolism | 0.3505926 | 0.02778041 |
| Cyanoamino acid metabolism | 0.350403 | 0.01341671 |
| Ether lipid metabolism | 0.346602 | 0.004607714 |
| p53 signaling pathway | 0.3424512 | 0.001598621 |
| Colorectal cancer | 0.3389656 | 0.002464333 |
| Small cell lung cancer | 0.3389656 | 0.002464333 |
| Toxoplasmosis | 0.3389656 | 0.002464333 |
| Viral myocarditis | 0.3389656 | 0.002464333 |
| Flavone and flavonol biosynthesis | 0.3362552 | 0.03059393 |
| D-Alanine metabolism | 0.3099026 | 0.04758065 |
| Flagellar assembly | -0.4501591 | 0.01617119 |
| Restriction enzyme | -0.4623062 | 6.60E-05 |
| Bacterial chemotaxis | -0.472489 | 0.006819423 |
| Bacterial invasion of epithelial cells | -0.5033562 | 2.75E-08 |
| Calcium signaling pathway | -0.5058255 | 1.74E-10 |
| Photosynthesis - antenna proteins | -0.5096998 | 1.60E-10 |
| Dioxin degradation | -0.5236634 | 0.000565156 |
| Chloroalkane and chloroalkene degradation | -0.5241747 | 0.000413393 |
| Bacterial motility proteins | -0.5446488 | 0.000532686 |
| Replication, recombination and repair proteins | -0.5446959 | 0.000426365 |
| Tetracycline biosynthesis | -0.5466325 | 0.000769446 |
| Xylene degradation | -0.6628035 | 2.05E-07 |
| Mineral absorption | -0.6630577 | 4.53E-09 |
| Sporulation | -0.7151384 | 1.23E-09 |
| African trypanosomiasis | -0.7423802 | 1.42E-10 |
| Germination | -0.7761985 | 6.15E-11 |
| Atrazine degradation | -0.8109147 | 2.62E-10 |
| Cytoskeleton proteins | -0.824936 | 1.86E-10 |
| Lipid metabolism | -0.8911117 | 1.40E-11 |

**Supplemental Table 3:**
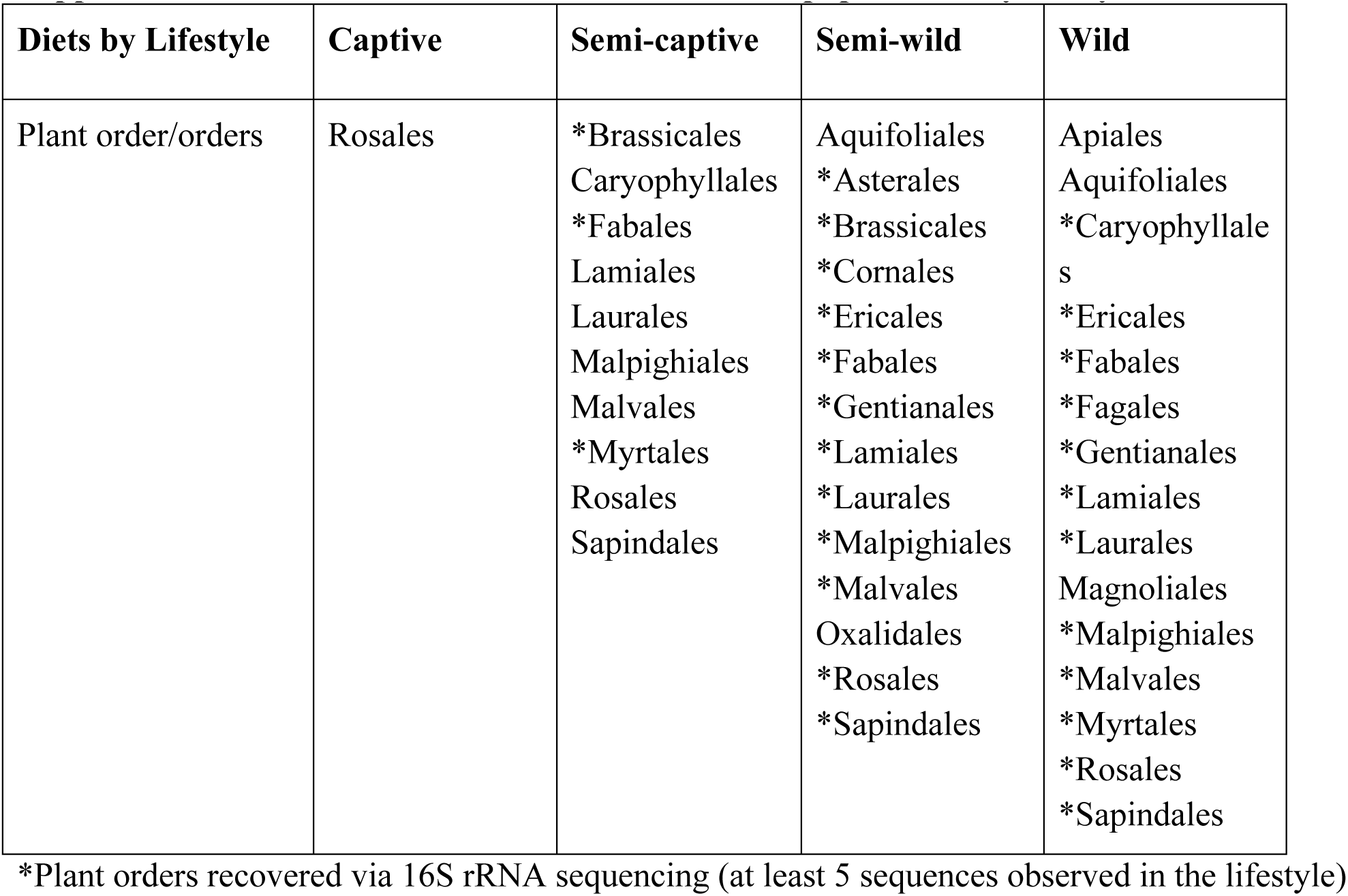
Plant orders observed in douc populations by lifestyle.

**Supplemental Figure 1.**
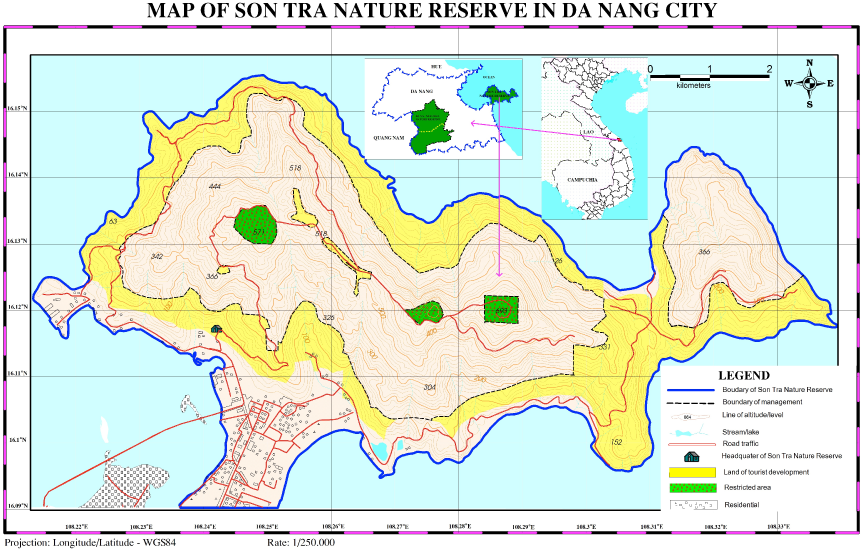
Study site. Red-shanked doucs (*Pygathrix nemaeus*) inhabiting Son Tra Nature Reserve, Da Nang, Vietnam (16°06’—16°09’N, 108°13’—108°21’E) served as the wild population for this comparative study (Lippold and Thanh 2008, Ulibarri 2013). Son Tra is located only 10 km from the heart of Da Nang City, which is the third largest city in Vietnam. The nature reserve is comprised of 4,439 total ha and of those 4,190 ha is covered by both primary and secondary forests (Lippold and Thanh 2008). Our study area was approximately 600 ha and is located on the north central region of the peninsula.

**Supplemental Figure 2.**
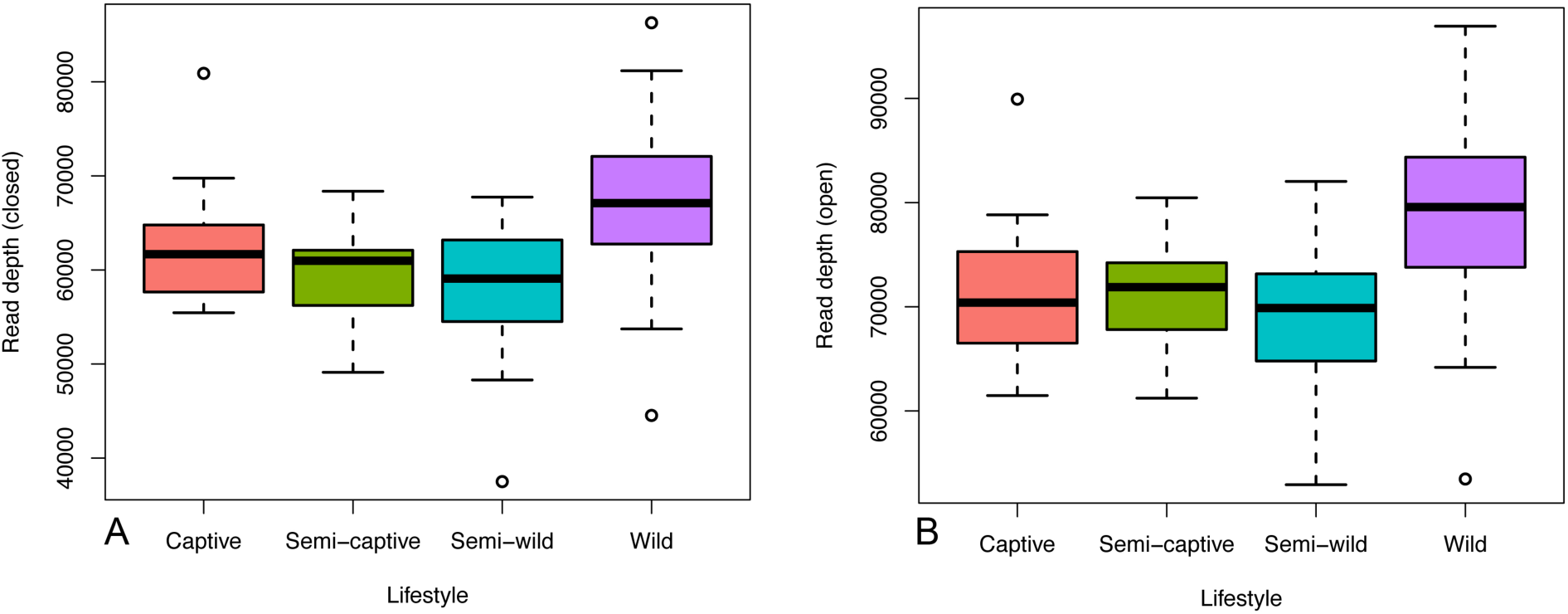
Read depth following quality control and OTU picking in the (a) closed-reference and (b) open-reference protocols. Read depth was relatively uniform across lifestyles in both protocols and highest in the wild population.

**Supplemental Figure 3.**
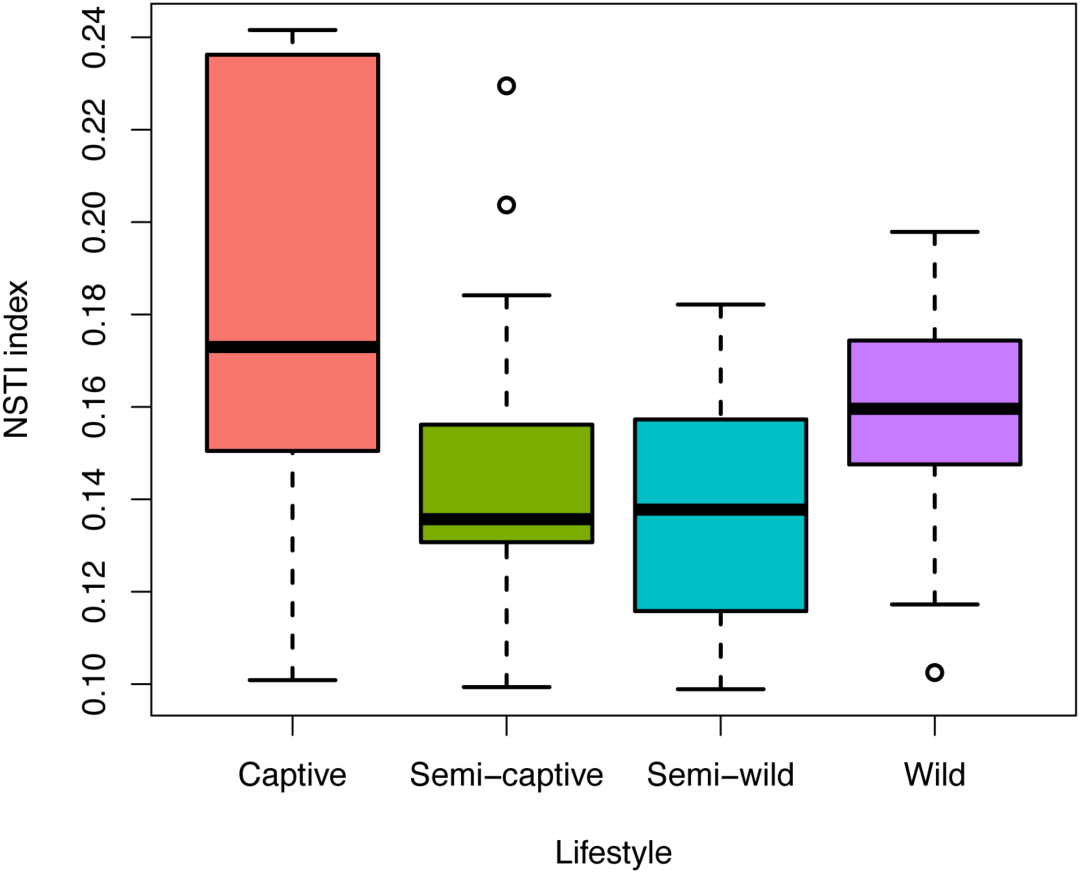
The Nearest Sequenced Taxon Index (NSTI) reported by PICRUSt for the lifestyle groups. The mean NSTI index for all groups was below 0.18, and below 0.165 for all lifestyles except captive. A lower NSTI index means less phylogenetic distance between the reference genomes used for functional predictions. A value of 0.17 was reported (Langille et al. 2013) to be a “medium” level of distance, indicating reasonable reliability for the PICRUSt functional predictions. Interestingly, the captive group showed the highest overall NSTI scores, potentially indicating a smaller proportion of genomic content in this lifestyle group has been fully sequenced.

**Supplemental Figure 4.**
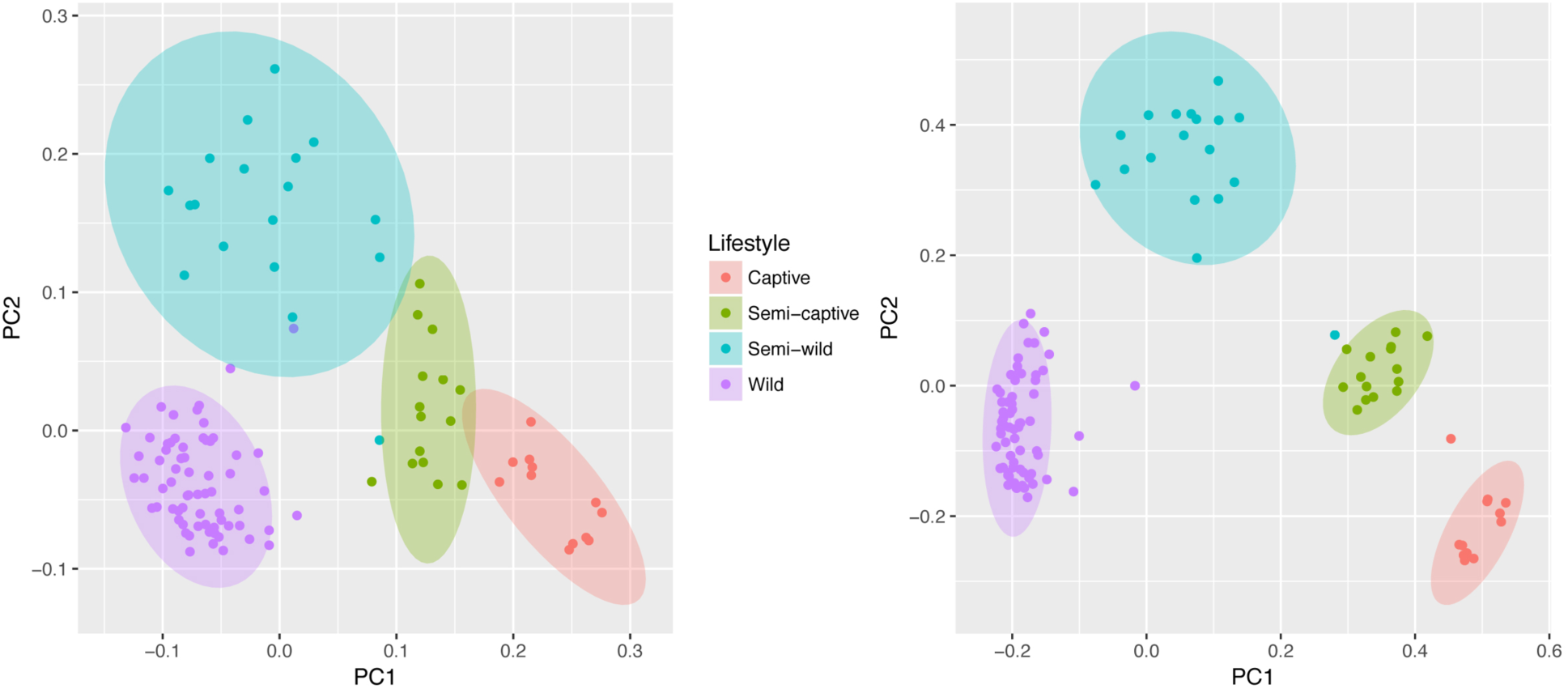
Principal coordinates plot of (a) weighted UniFrac and (b) Bray-Curtis metrics showing ecological distance between gut microbial communities in wild, semi-wild, semi-captive, and captive red-shanked doucs. All samples were obtained with the same protocol for V4 16S rRNA sequencing, and open-reference OTU picking was used. Douc microbiomes clearly clustered by population suggesting that each douc population had a unique microbiome, and thus were highly distinctive from one another.

**Supplemental Figure 5.**
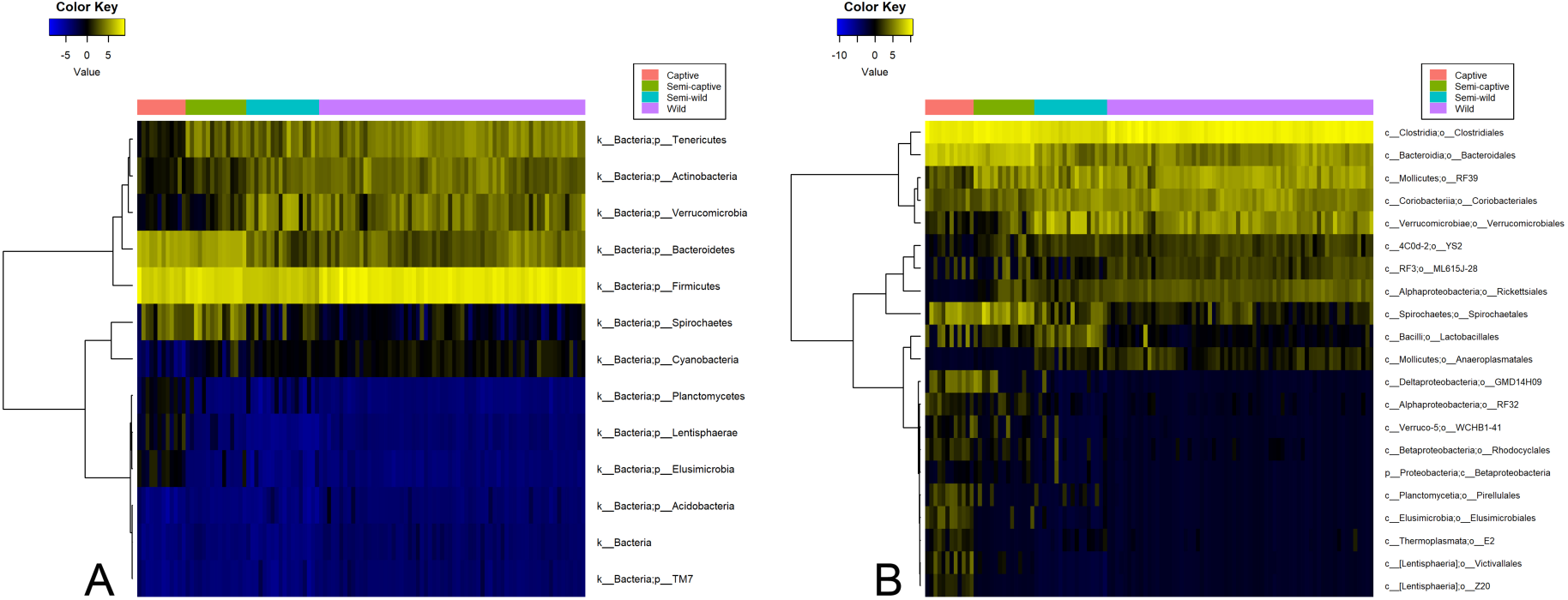
Heatmaps of differentially abundant microbial taxa at the phylum and order levels in red-shanked doucs living four distinct lifestyles. Taxa are displayed with polyserial correlations (rho) above 0.3, rho estimate adjusted p < 0.05, and (pairwise) Wilcoxon rank-sum adjusted p-value for wild and captive lifestyles < 0.05. Color represents intensity of centered log ratio abundances along gradient of color scale shown, at a) phylum and b) order levels of taxonomy.

**Supplemental Figure 6.**
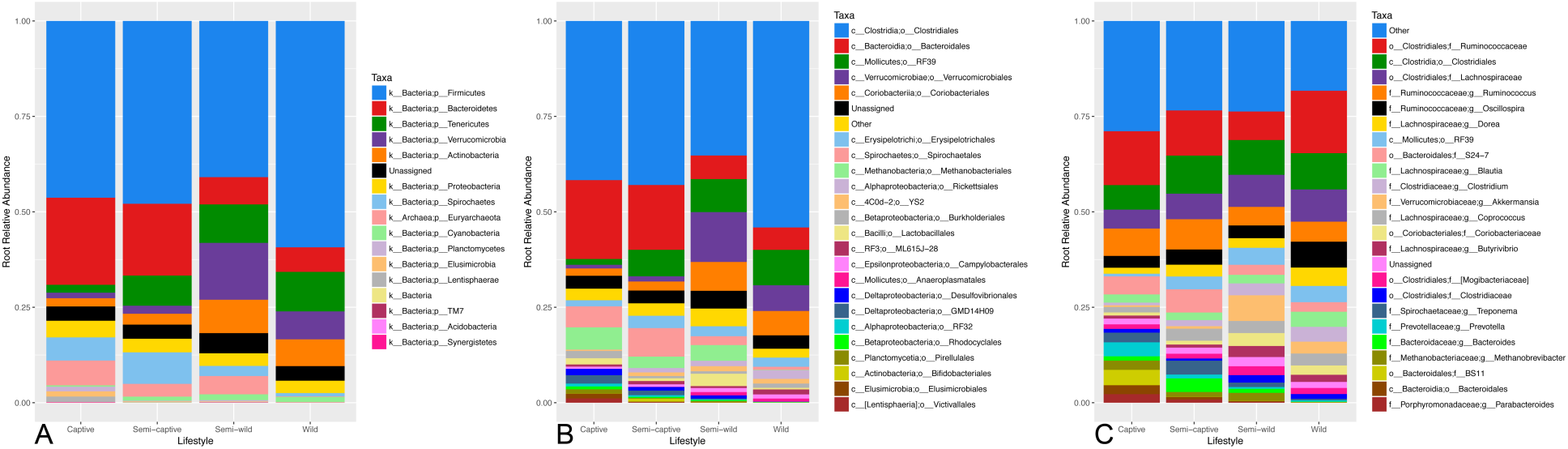
Stacked bar plots of microbial taxa relative abundance ordered by taxonomic level (phylum, order, and genus) in wild, semi-wild, semi-captive, and captive red-shanked doucs. All samples were obtained with the same open-reference V4 16S rRNA protocol. Up to 25 taxa (including a group for “Other”) are displayed. Taxa were prioritized for display according to highest groupwise maximum abundance, and ordered for display by average abundance throughout the dataset from top to bottom. For display purposes, the square root transformation of the abundance is shown rather than the absolute relative abundance to allow for easier visual inspection and comparison of lower-abundance community members. Legends show the known taxonomic rank of each taxon in the graph; colors may not be consistent between graphs. Microbes that could not be classified to the specified level are also included at the highest level at which they could be classified.

**Supplemental Figure 7.**
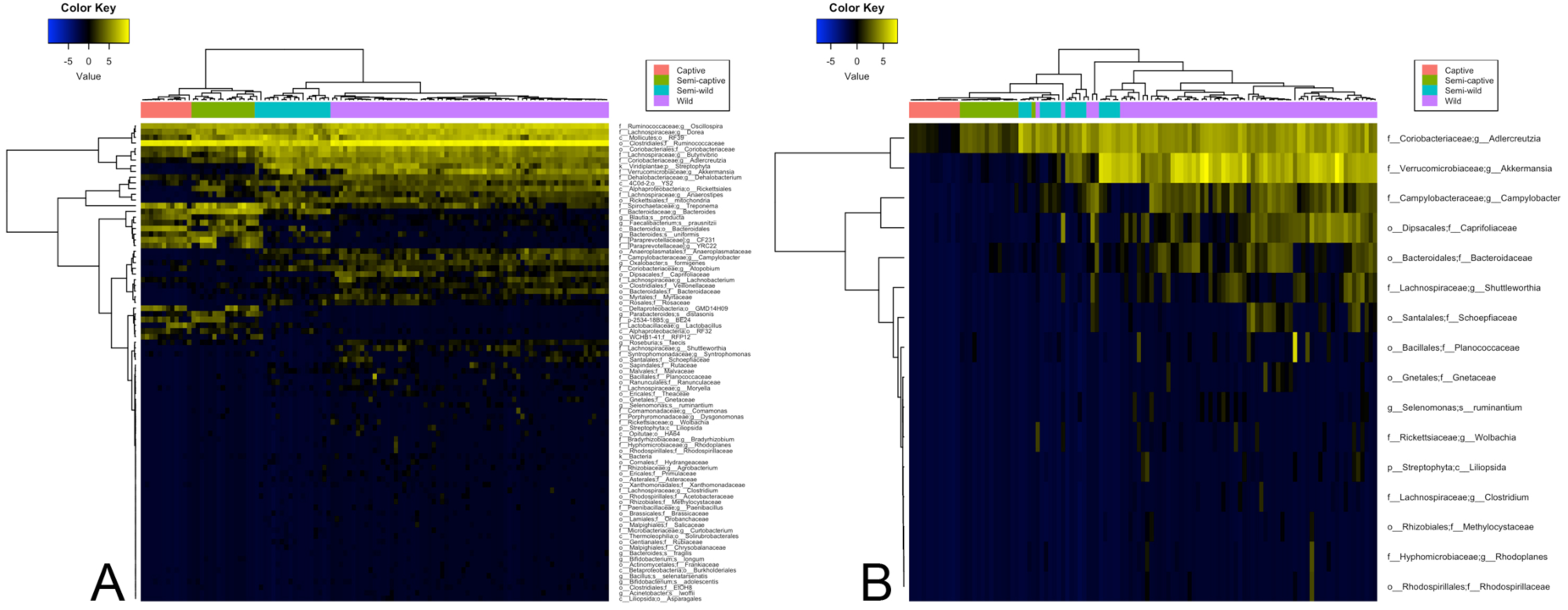
Biclustering of significantly differentiated taxa recapitulates group separation by lifestyle. Heatmaps at different levels of significance were generated using all of the statistically significant finest-grain features available in the data (up to the order level in plants and species level in prokaryotes). Biclustered heatmaps were generated with the following taxa selection criteria: (a) polyserial rho > 0.45 and captive-wild adj. Wilcoxon p < 0.01 (84 taxa selected); and (b) polyserial rho > 0.75 and Wilcoxon p < 0.001 (16 taxa selected). Both heatmaps were subjected to unsupervised complete linkage clustering of correlation dissimilarity [1 - cor()], revealing that the correlation patterns of top differentiated taxa alone is sufficient to nearly completely recover lifestyle group membership of the samples. Further, increasing stringency of the selection criteria as in (b) resulted in limited blurred membership between adjacent lifestyle wildness categories only. This highlights the potential utility of these taxa as biomarkers in this population, as well as efficacy of the gradient-informed criteria used in their selection.

**Supplemental Figure 8.**
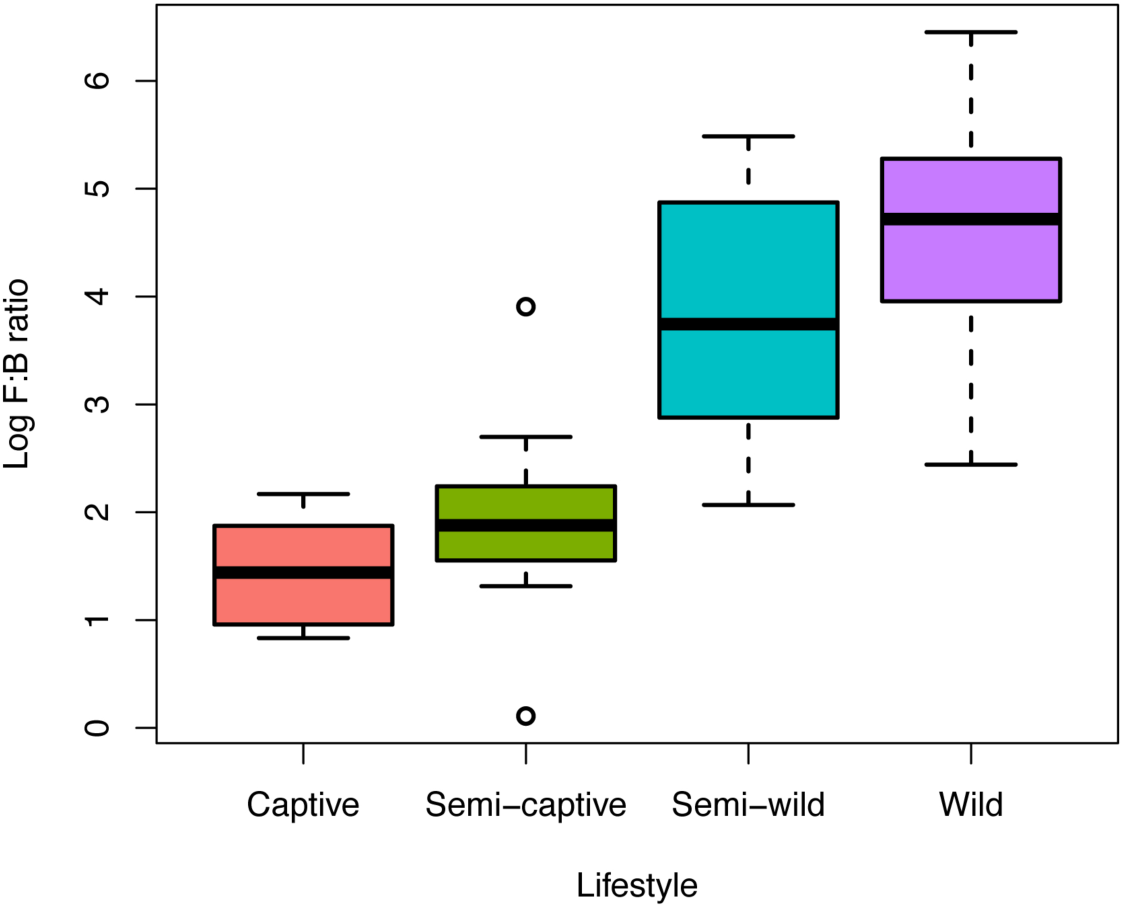
Firmicutes to Bacteroidetes ratio in red-shanked douc microbiomes across populations. Bar plots of *Firmicutes* to *Bacteroidetes* ratio, a measure of energy harvest capacity by microbial communities, plotted by wild, semi-wild, semi-captive, and captive populations of red-shanked doucs. All samples were obtained with the same protocol for V4 16S rRNA sequencing, and open-reference OTU picking was used. The *Firmicutes* to *Bacteroidetes* ratio was highest in wild doucs (4.64 ± 0.94), followed by the semi-wild doucs (3.78 ± 1.14), semi-captive (1.94 ± 0.81), and captive doucs (1.43 ± 0.50). A significant decrease in *Firmicutes* to *Bacteroidetes* ratio from the wild population to the semi-wild population, from semi-wild to the semi-captive population, and again from the semi-captive population to the captive population is visible. Pairwise Wilcoxon rank-sum p-values for the populations are: captive vs semi-captive p = 0.052858, captive vs semi-wild p = 4.62464e-08, captive vs wild p = 4.320356e-08, semi-captive vs semi-wild p = 5.177609e-06, semi-captive vs wild p = 6.900029e-09, semi-wild vs wild p = 0.006974937.

